# *Least* Component Analysis reveals ecological constraints in microbial communities

**DOI:** 10.64898/2026.05.22.727221

**Authors:** Víctor Peris-Yagüe, Stella Perez, Daniel R. Amor, Simona Cocco, Rémi Monasson

## Abstract

How microbial communities maintain robust and reproducible ecological functions despite their extraordinary taxonomic diversity remains an open question. Here we show that functional organization in microbial communities can be uncovered by repurposing Principal Component Analysis to focus on directions of lowest variance in taxon abundance data, rather than maximal variance. These least-variance components are statistically significant and correspond to ecological constraints on taxon abundances that are consistently fulfilled across samples. Using consumer-resource models, we show that these constraints arise from resource-mediated interactions and express biomass conservation, effectively grouping taxa into producer and consumer guilds. We validate this interpretation in simulated communities and experimental systems under competition and cross-feeding. Finally, we show that low-variance structure is ubiquitous in natural microbial communities and reveals a sparse network of taxa with disproportionate influence on community structure. Together, our results establish low-variance components as indicators of ecological constraints linking taxonomic diversity to functional organization.

## 1 Introduction

Microbial communities harbor extraordinary taxonomic diversity while performing robust and reproducible ecological functions [1]. Surveys spanning thousands of environments—from ocean surface waters, to soils and the human gut —reveal communities composed of hundreds to thousands of co-occurring taxa, whose identities are shaped by geography and environmental selection [1, 2, 3]. Yet beneath this taxonomic variation, community-level functional profiles remain comparatively stable: distinct groups of taxa reproducibly perform the same metabolic roles across ecosystems, a pattern often attributed to widespread functional redundancy [4]. Interpreting this apparent decoupling between taxonomic and functional variation remains challenging, as reduced functional variability can arise from statistical averaging rather than from active selection on community traits [5, 6]. This raises a fundamental question: how is functional organization maintained in systems composed of hundreds of interacting taxa?

Ecological interactions among community members, including competition for resources [7] and facilitation—including metabolic cross-feeding [8, 9]—can have a significant effect on the functional robustness of microbial communities[10, 11]. Cross-feeding allows multiple taxa to coexist in communities supplied with a single resource [12, 13], as metabolic by-products released by one taxon can serve as resources for others. A generic and widely observed example is the interaction between fermentor and respirator taxa, in which fast-growing fermentors secrete organic acids during growth on sugars—a consequence of overflow metabolism [14]—which in turn support the growth of respirators specialized on these metabolites [15]. Such interactions can give rise to reproducible metabolic organization even when taxonomic composition varies across communities [12]. While these interactions explain how functional structure can emerge from diverse communities, they do not directly specify how taxon abundances are quantitatively constrained. This gap is particularly striking given that most available data consist primarily of abundance measurements across samples.

In response to this limitation, a range of quantitative approaches has been developed to infer microbial interactions and functional organization directly from abundance data [16, 17]. Several methods aim to identify groups of taxa whose collective abundances best predict observed community functions, commonly referred to as functional guilds, even in the absence of explicit mechanistic understanding [18, 19, 6]. More generally, dimensionality-reduction techniques such as principal component analysis (PCA)[20] are routinely applied to microbial abundance datasets to uncover low-dimensional structure. For example, PCA has revealed metabolically coherent groupings in fluctuating environments [21]. However, because PCA emphasizes directions of maximal variance across samples, it inherently overlooks structure encoded in low-variance directions, which are typically relegated to noise.

Inspired by protein sequence analysis—where low-variance directions capture functional constraints rather than noise [22]—we propose extending this perspective to microbial ecology. Here, we show that directions of lowest variance in PCA, the Least Components (LCs), encode statistically significant structure in microbial communities that is directly linked to their functional organization. In microbial abundance data, LCs encode conservations of biomass that emerge from resource-mediated interactions under both competition and cross-feeding. We show that these low variance constraints group taxa into metabolic guilds, such as producers and consumers, in experimental communities, and that LCs are prevalent in natural communities. Together, our results establish directions of low variance as a quantitative link between taxonomic diversity and functional organi-zation, and identify least components of PCA as a simple yet powerful tool for uncovering ecological constraints from abundance data.

## 2 Results

### 2.1 Least Components identify directions of restricted variation in taxon abundances

Microbial community datasets often consist of absolute or relative abundances of different taxa across multiple samples (Fig. 1a). Each sample therefore corresponds to a point in a high-dimensional space of species abundances (orange dots in Fig. 1b). PCA is routinely used to uncover structure in the distribution of points by focusing on directions with highest spread, the Principal Components (PCs; green arrow in Fig. 1b).

**Figure 1.**
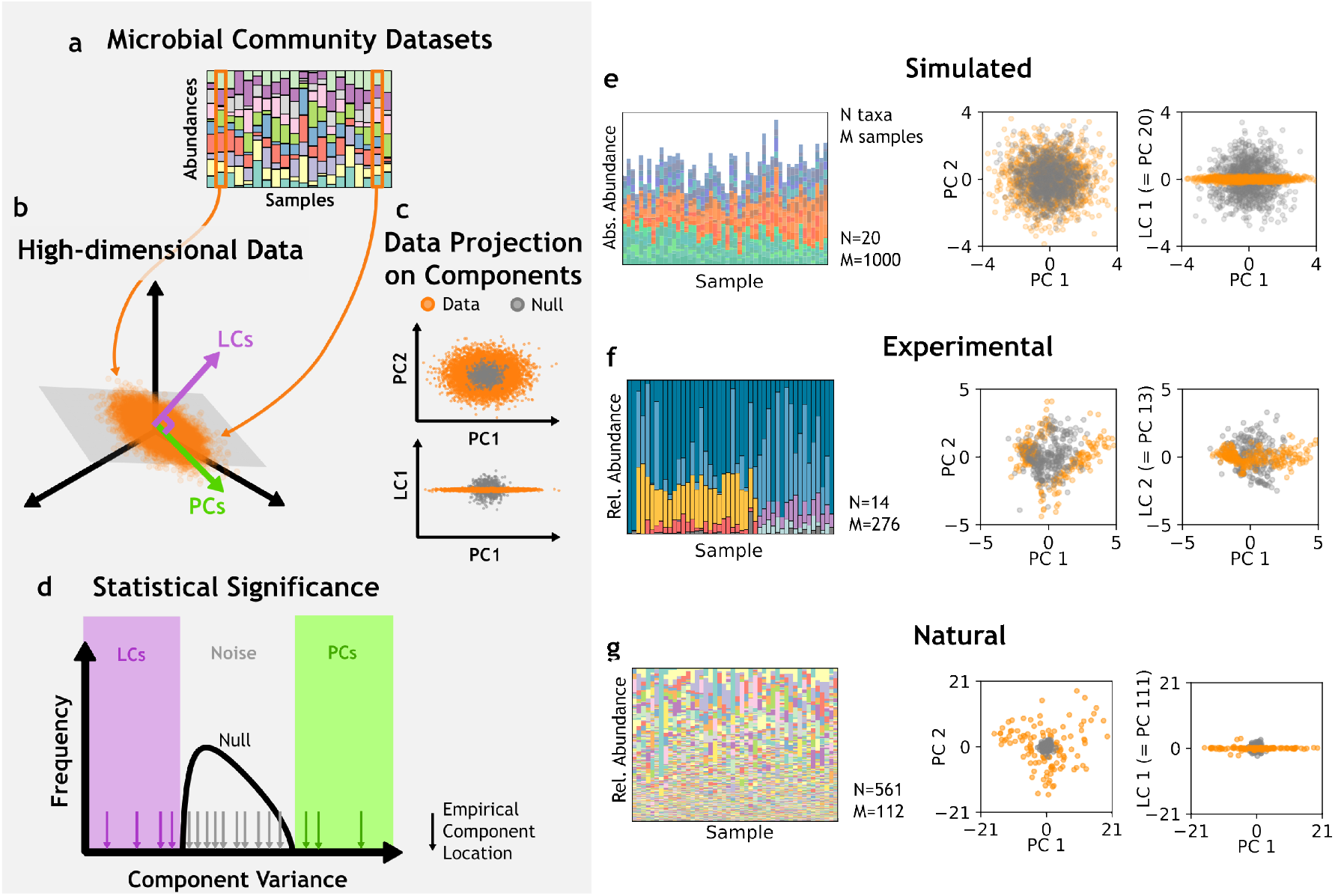
Principal and Least Components identify directions of, respectively, large and low variance across ecological datasets. **(a)** Schematics of species abundances (colors) in representative samples of a microbial community dataset. **(b)** The distribution of samples (orange dots) in abundance space (axes) exhibits directions of large variance, captured by PCs (green), and low variance, captured by LCs (purple). **(c)** Projections of empirical data along PCs (LCs)—component variance—exhibit larger (smaller) variance than expected under a structureless null model. **(d)** Component variance is deemed statistically significant when incompatible with the appropriate null model. Components whose variance fall within the null expectations are attributed to noise. **(e-g)** Taxon abundances (left) in representative samples, as well as their projections on PC1-PC2 (center) and PC1-LC1 (right), for a simulated community under a Consumer-Resource framework with cross-feeding (e), an experimental [15] community of soil origin (f), and a natural [23, 24] soil community (g). Colors in barplots, individual taxon; orange dots, empirical samples; gray dots, null-model samples.

Although typically overlooked, directions with smallest spread, the Least Components (LCs; purple arrow in Fig. 1b) are also indicators of non-trivial structure[22]. Ecological structure, for example, causes sample projections onto principal and least components to exhibit, respectively, larger or smaller spread than expected from a null model with the same numbers of samples and taxa but no structure (Fig 1c). Component variances—the variance in the data projected on a component— that fall beyond the upper or lower tails of the null distribution define statistically significant PCs and LCs, respectively (Fig. 1d, Methods). Importantly, LCs can contribute on equal footing with PCs to explaining data structure (in the context of Maximum Likelihood inference; SI Section SI 1)[25].

To assess the existence and prevalence of LCs in microbial communities, we examined datasets for three different microbial communities (Fig. 1e-g,left). The first dataset consists of community samples simulated via Consumer-Resource dynamics until resource depletion in a closed system, using random initial values for resource concentrations and species abundances (Methods). The second dataset comprises final taxon abundances from a serial dilution experiment in which replicate cultures were inoculated from a single soil inoculum [15]. The third dataset consists of soil samples from an agriculture study of wilt disease in tomato plants [24].

After rescaling taxon abundances to z-scores, we performed PCA by diagonalizing the empirical Pearson correlation matrix of each dataset (Methods). Figure 1e-f shows that, for the three communities, data points display larger spread when projected along the top PCs than the corresponding null data (Methods). All three datasets have statistically significant LCs, where data projections display lower variability than the null model. These LCs constitute linear combinations of taxon abundances that remain tightly conserved across samples, reflecting the presence of strong ecological constraints within each community.

### 2.2 Least Components reveal biomass conservation emerging from resource competition in microbial co-cultures

To explore how least components can emerge from ecological constraints, we first consider a minimal community dominated by a canonical ecological interaction: competition for a single limiting resource. Specifically, we study two closely related *Escherichia coli* K-12–derived strains carrying distinct markers—a reference strain (E_*K*_) and a fluorescently labeled strain (E_*F*_)—growing in a glucose-limited medium. When separately characterized in monoculture, E_*K*_ and E_*F*_ have similar yields—biomass produced per unit of glucose consumed—with the latter strain exhibiting a higher growth rate (Fig. 2a, left).

**Figure 2.**
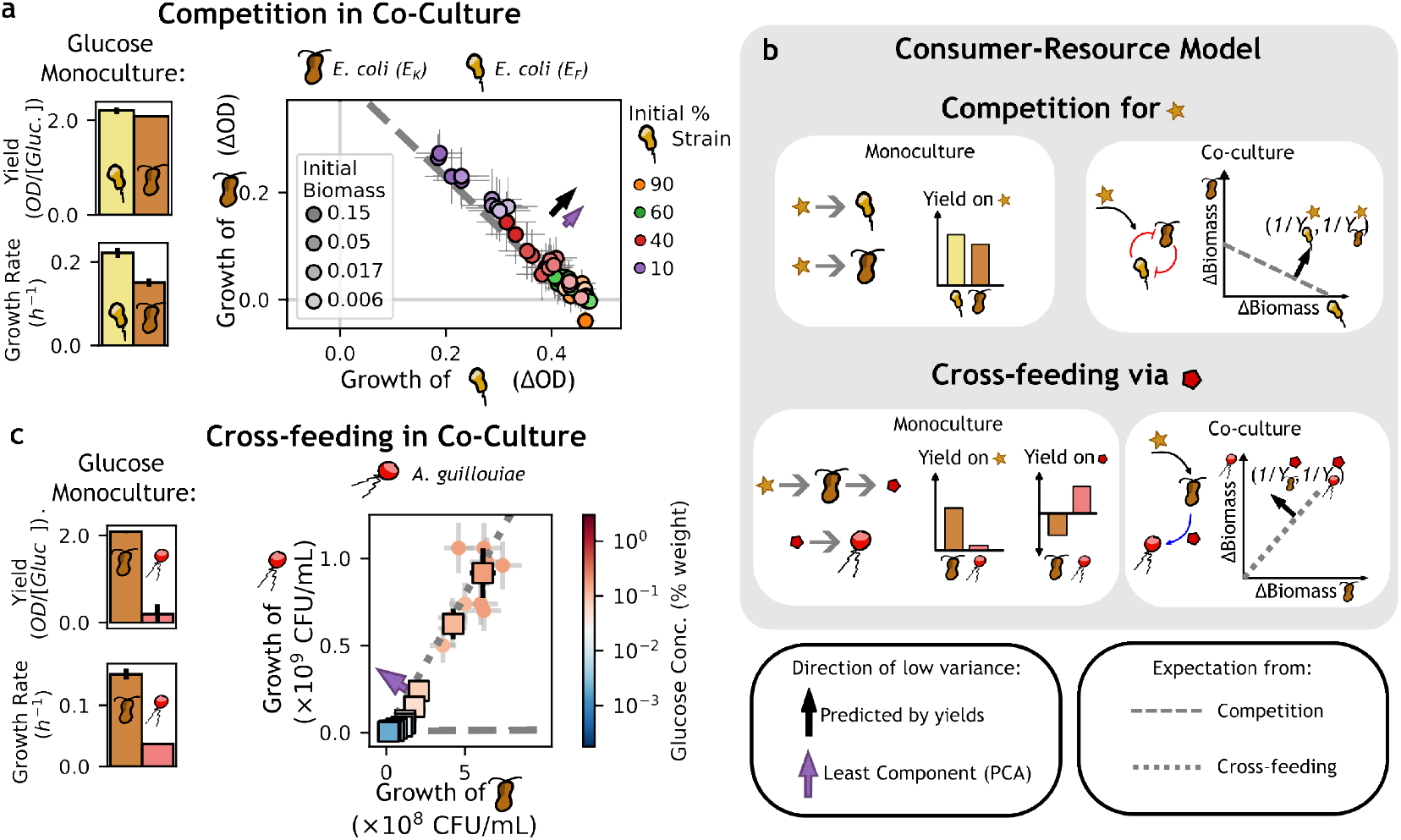
Resource-mediated conservation of biomass leads to different directions of low variance under competition and cross-feeding in bacterial co-cultures. **(a)** Left: Monoculture yields (top) and growth rates (bottom) of two *E. coli* strains—a non-fluorescent strain E_*K*_, in brown, and a fluorescently labeled strain E_*F*_, in yellow—grown on glucose as the only carbon source. Right: growth of each strain from inoculation to saturation in co-culture (Methods). Dots stand for individual replicates (with a minimum of 2 per condition), color indicates initial percentage of E_*F*_ biomass, dot transparency indicates initial co-culture biomass, error bars refer to fluorescence-based biomass inference (Methods). **(b)** Consumption and production yields define directions of low variance of species abundances in consumer-resource models (Eq. (1)). Under resource competition (top), the entries on the vector of inverse yields have the same sign. Under cross-feeding (bottom), they have opposite sign. **(c)** Composition in a two-species cross-feeding experiment between E_*K*_ and *A. guillouiae* (A_*g*_) strain in minimal media supplemented with different glucose concentrations (color bar). Dots correspond to individual replicates, errorbars correspond to standard deviation from CFU (Methods). Squares stand for averages over a minimum of 4 replicates for each glucose concentration (color). **All panels:** Black (resp. purple) arrows mark the direction of inverse yields (resp. LC). The length of the purple arrows in panel (a,c) is 5 × the standard deviation of the data in its direction. A dashed (resp. dotted) gray line is the expectation under competition (resp. cross-feeding).

Measurements of strain growth in batch co-cultures initiated from a range of initial abundances (Methods and Fig. SI 10) reveal that final abundances cluster along a line (Fig. 2a, right). Data show substantial variability along this line and are biased toward higher growth of E_*F*_, consistent with its higher growth rate in monoculture. The linear relationship between abundances results from the competition for a limiting resource, in which the growth of one strain constrains that of the other. PCA of the final abundances identifies a LC (purple arrow in Fig. 2a; Methods), approximately perpendicular to this line.

The consumer-resource (CR) framework[26, 27] provides a mechanistic explanation for how compe-tition for a resource gives rise to a LC. In a CR model, each taxon (or strain) *i* grows in abundance by consuming a resource α, with growth proportional to the amount of resource taken up. Under the assumption of constant yields 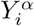, conservation of biomass from the shared resource imposes constraints on the final taxon abundances that can be expressed as

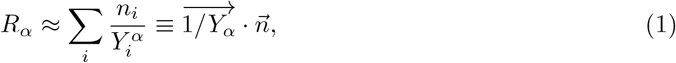

where R_*α*_ ≥0 denotes the amount of resource α externally supplied to the system. Here, *n*_*i*_ is the final abundance of taxon *i*, which approximates the net change in abundance once the supplied resource has been depleted (Methods). This resource-mediated constraint expresses a conservation of biomass at the community level: the total biomass that can be produced is limited by available resources, imposing linear restrictions on the space of possible community compositions, analogous to conservation laws in physics and chemistry. As a result, abundances are free to fluctuate within a hyperplane—which in the case of two species is simply a line—and are tightly constrained along the orthogonal directions (Fig. 2b), defined by the inverse-yield vectors.

This theoretical prediction aligns closely with our experimental findings for two *E. coli* strains competing for glucose in Fig. 2a: the direction along the LC has slope ≈ 1.012, in good agreement with the ratio of inverse yields measured in monocultures, 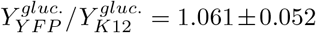 (Methods). Fluctuations in the experimental system give rise to the small variance of the data around the linear constraint. This variance is quantified by the eigenvalue associated with the LC (Fig. 1a,b) and is predicted to increase with fluctuations in both resource and species dynamics (Fig. SI 11 and Fig. SI 2).

### 2.3 Least Components capture biomass conservation under cross-feeding

The derivation of the CR conservation relation in Eq. (1) also extends to the case of negative yields 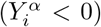, which can be interpreted as net production of a resource by a taxon. Such secondary resource may then support the growth of another taxon through a cross-feeding interaction. We therefore expect the growth of the two taxa to be proportional. The resulting line of positive slope remains orthogonal to the inverse-yield vector associated with the cross-fed resource (Fig. 2b, bottom), which is not externally supplied (*R*_*α*_ = 0).

To experimentally test this prediction, we co-cultured *E*_*K*_ with an *Acinetobacter guillouiae* (*A*_*g*_) strain that grows poorly on glucose but exhibits strong growth on organic acids (Fig. 2c and Fig. SI 12). This pairing provides a model system for a common fermentor–respirator interaction [12], in which sugar-utilizing taxa produce organic acids that support the growth of specialist consumers. We find that *A*_*g*_ grows robustly in co-culture with *E*_*K*_ over a range of initial glucose concentrations, despite glucose being the only externally supplied resource (Fig. 2c and Fig. SI 13). PCA of the co-culture data reveals a LC (purple arrow in Fig. 2c), along which abundances are tightly constrained. The slope of the LC direction gives indirect access, according to the CR model,to the ratio of the yields of the two strains on acid secretions, 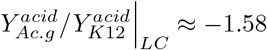.

Together, these results suggest that LCs can capture resource-mediated constraints operating in competitive as well as in cross-feeding microbial communities. We stress that these findings are not restricted to closed systems in batch culture, but also generalize to CR models for open systems with resource influx and biomass outflow (SI Section SI 2), such as chemostats.

### 2.4 Least Components map to ecological interactions in simulated and experimental microbial communities

For communities with more than two taxa, the CR framework predicts linear constraints (Eq. (1)) that define hyperplanes in the space of taxon abundances, one for each resource. These constraints express a conservation of biomass at the community level, limiting the total biomass that can be produced from available resources. As a result, abundances are free to fluctuate inside a manifold defined by the intersection of the hyperplanes, but are tightly constrained along the orthogonal directions corresponding to the inverse-yield vectors. We tested these predictions by analyzing data from more complex microbial communities.

We first analyzed the enrichment experiment of Estrela *et al*.(Fig. 1f) [15], which provides a wellcharacterized example of microbial cooperation. Communities were grown with glucose as the sole externally supplied carbon source, primarily fermented by two Klebsiella taxa (*Km* and *Kp*). Fermentation leads to the secretion of organic acids, which are subsequently consumed by two respiratory taxa, *Alcaligenes* and *Pseudomonas*, and an additional cooperative interaction between *Alcaligenes* and a *Delftia* strain has been reported [15] (Fig. 3a).

**Figure 3.**
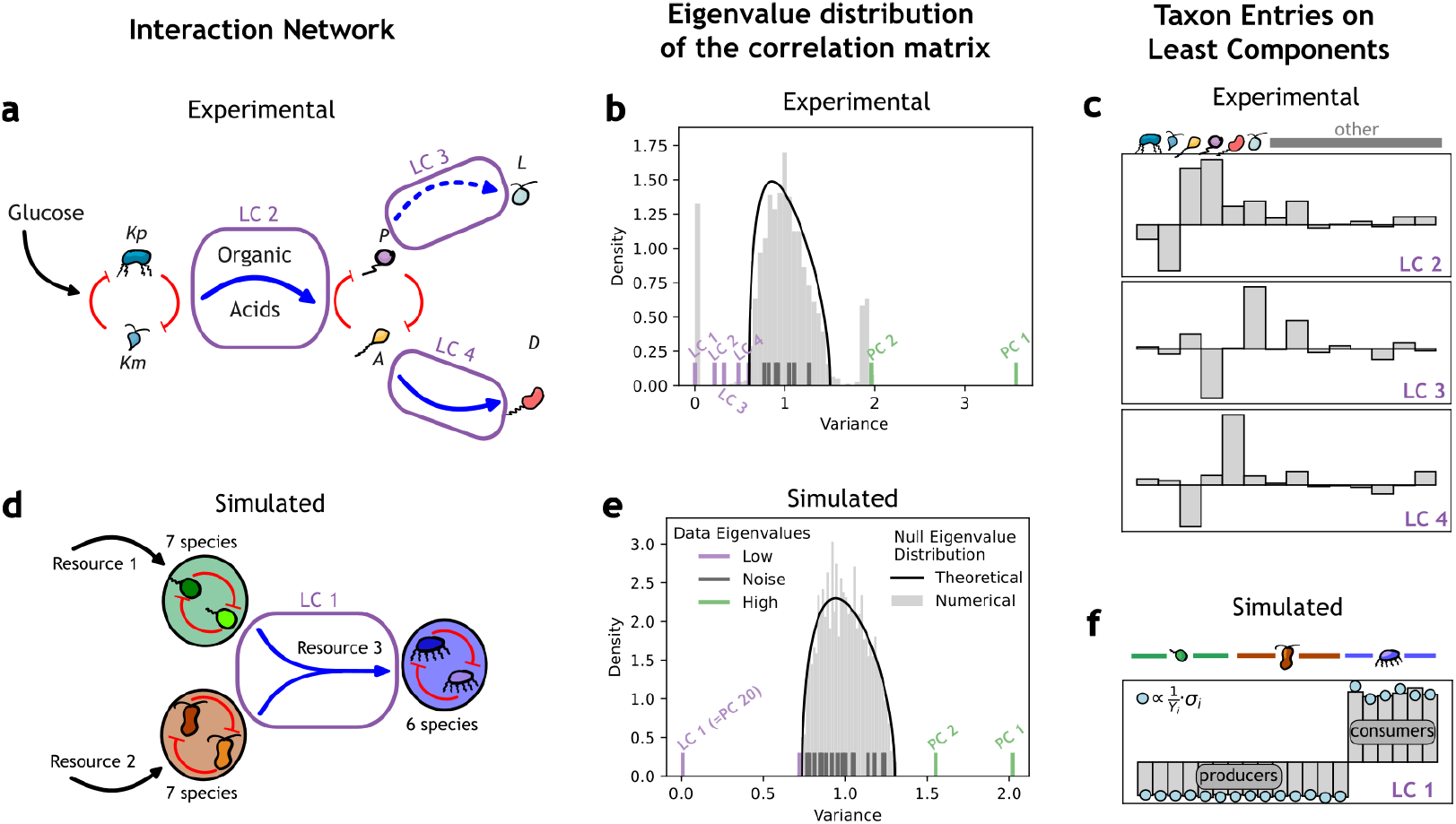
Least components infer resource-mediated interactions in microbial communities. **(a)** Resource-mediated interaction network of the experimental community. Black arrows indicate externally-supplied resources, red arrows indicate competition for a resource, blue arrows indicate facilitation, potentially mediated by cross-feeding (resources named on the arrows when known). Solid arrows denote previously reported interactions [15]; whereas dashed arrows denote novel interactions inferred by LCs. **(b)** Variance of components for the experimental dataset, compared with two null expectation: a theoretical null (black line; Methods) and a numerical null (grey histogram; Methods). **(c)** Individual species entries for LC2 (top), LC3 (center), and LC4 (bottom) of the experimental dataset. **(d, e)** As in (a) and (b), respectively, resource-mediated interaction network and variance of components for the simulated community. **(f)** LC 1 of the simulated dataset. Blue dots denote entries of inverse yields vector, properly rescaled (Methods).

Applying PCA to this dataset reveals two significant principal components (PC1 and PC2) and four least components (LC1-LC4) (LCs are numbered in increasing component variance; Fig. 3b). PC1 separates samples according to the dominating respirator strain (Fig. 1f and Fig. SI 14). LC1 and PC2 arise from the relative abundance nature of the dataset and do not capture ecological structure, as confirmed by a null model adapted to relative abundances (Fig. 3b, Methods, SI Section SI 4).

The remaining three least components (LC2-LC4) reflect distinct interactions in the community. LC2 captures the cross-feeding of organic acids: the component entries of *Kp* and *Km* share one sign, whereas those of *Alcaligenes* and *Pseudomonas* have the opposite sign, consistent with producer-consumer roles. Although LCs are defined up to a global sign, prior experimental characterization [15] allows for the unambiguous identification of *Klebsiella* taxa as producers and *Alcaligenes* and *Pseudomonas* as consumers. Likewise, LC4 similarly captures the facilitation between *Alcaligenes* and *Delftia*. LC3 infers a cooperative interaction between *Pseudomonas* and a *Lachnoclostridium* taxon (dashed blue arrow in Fig. 3a, which was not reported in [15] but it is supported by co-occurrence statistics; p_*val*_ < 10^*−*15^; Methods). The smaller eigenvalue associated with LC2 compared with LC3 and LC4 suggests that abundance fluctuations increase further downstream from the externally-supplied resource, indicating that fluctuations accumulate along cross-feeding interactions. To test the generality of our findings, we analyzed an independent enrichment dataset [12] and again found that least components capture the generic cross-feeding interaction between fermentor and respirator taxa reported in that system [12] (SI Section SI 5).

We next analyzed the simulated microbial community dataset (Fig. 1e). In this system, species consume one of three resources, defining three groups (green, orange, blue in Fig. 3d; Methods). Two resources are externally supplied in variable amounts across samples, whereas a third resource is produced internally by species in the community. In this case, PC1 separates species according to their preferences for the externally supplied resources, while PC2 encodes overall abundance fluctuations (Fig. SI 15). In contrast, LC1, the third and last significant component of this system, reflects interactions mediated by the internally generated resource (Fig. 3f): consumer species (blue) share one sign in their component entries, whereas producer species (green and orange) share the opposite sign. Beyond sign structure, the magnitudes of the entries quantitatively agree with the inverse yields of species on the internally generated resource, as predicted by the CR model (Eq. 1). Similar results are obtained for a community with identical structure, simulated under chemostat conditions (Fig SI 16).

Across both datasets, least components robustly reveal facilitation but do not systematically capture competition for externally supplied resources. In the simulated dataset, large sample-to-sample variability in the supply of resources (*R*_*α*_, α ∈ {1, 2} in Eq. 1) gives rise to the two principal components. However, reducing variability in the supply of one resource gives rise to a second least component that encodes competition among its consumers (Fig. SI 17). In the experimental dataset, although the supplied resource amount was held constant across samples, the information on competition is lost due to the relative abundance nature of the data. Eq. 1 holds for absolute taxon abundances associated with a resource supplied at constant amount. When considering relative abundances, the resource (left term of Eq. 1) must be rescaled by the total sample abundance, which fluctuates from sample to sample (Fig. SI 17 and SI Section SI 6). Facilitation, by contrast, corresponds to a constraint on internally generated resources (*R*_*α*_ = 0) and is therefore invariant under the rescaling. Restrictions from facilitation can therefore be fulfilled even in fluctuating environments (Fig. SI 17b) and are more robust to distortions due to measuring relative, rather than absolute, abundances (Fig. SI 17c).

### 2.5 Projections onto least components capture functional activity under facilitation

We examined how community samples fulfill the abundance constraints encoded in least components through different community compositions, focusing on cross-feeding. Under the consumer-resource interpretation of least components, we grouped taxa according to their contribution on each LC: producers, corresponding to taxa with negative component entries, and consumers, corresponding to taxa with positive entries. We then define, for each sample, the total consumption 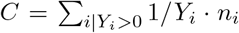and total production 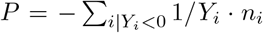 associated with the resource (Fig. 4a). At equilibrium, consumption and production are balanced in most samples, up to fluctuations quantified by the variance of the least component (Fig. 4a). Along this equilibrium line, the distance to the origin indicates overall functional activity—joint production and consumption of the resource.

**Figure 4.**
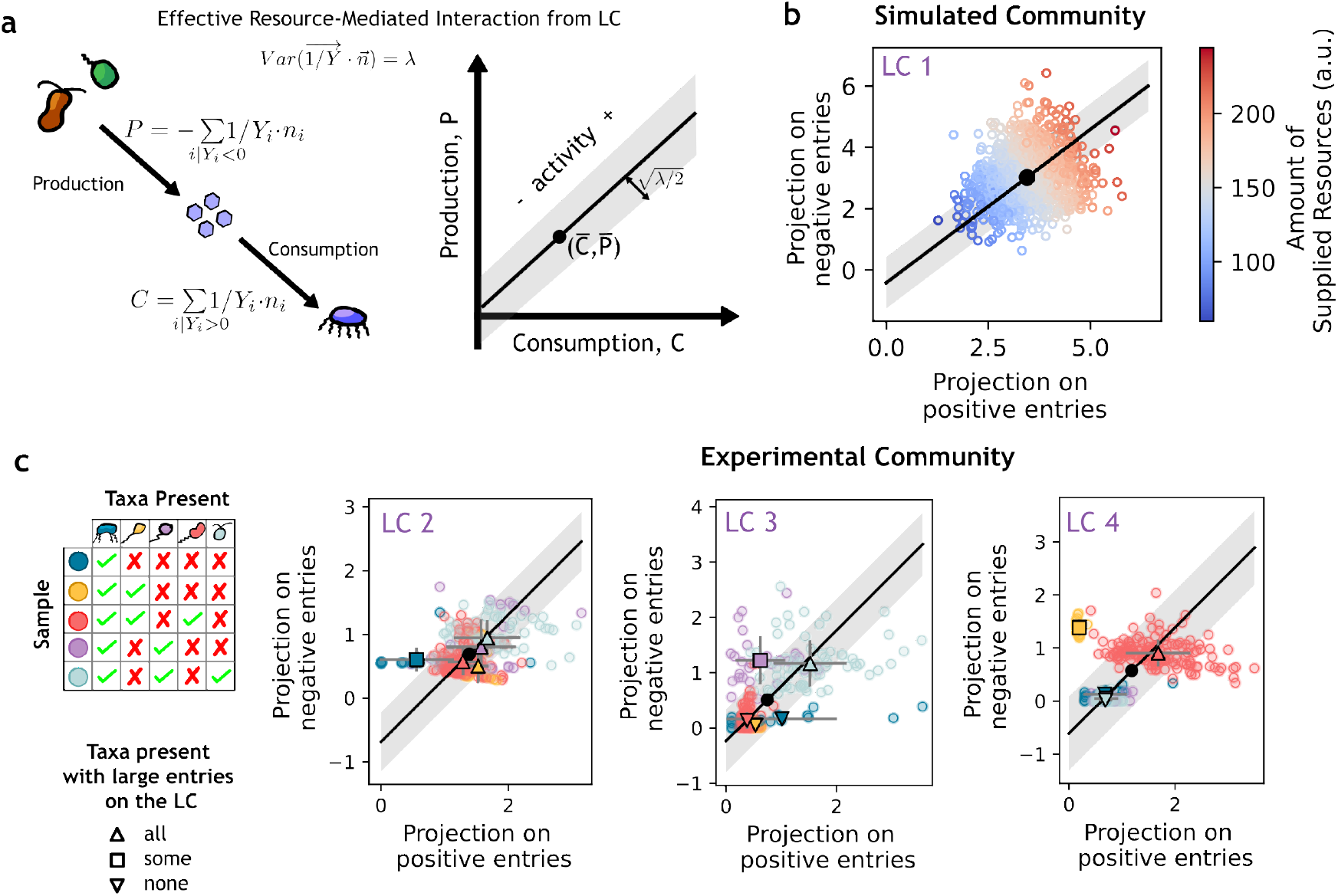
Projections onto least components encode functional activity and reveal extinction of key taxa. (**a)** Schematic interpretation of least-component projections under the Consumer-Resource framework. Positive and negative component entries correspond to consumer (C) and producter (P) contributions, respectively. Samples lie close to the equilibrium line C = P, with fluctuations set by the standard deviation of the least component, 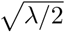. Distance from the origin indicates functional activity—joint production and consumption of the resource. **(b)** Projection of samples on the positive and negative entries of LC1 for the simulated community dataset. Functional activity scales with the amount of externally supplied resources (color scale). **(c)** Same analysis applied to the experimental community dataset [15]. Circles indicate individual samples, colored by taxonomic composition. Other symbols stand for averages over samples of the same composition (legend shown on the left). Triangles denote samples in which all taxa with large component entries on the LC are present, inverted triangles denote samples in which these taxa are absent, corresponding to extinction of functional taxa, and squares denote samples in which only a subset of these taxa is present. Panels show projections onto LC2, LC3, and LC4 from left to right.

We first applied this analysis to the simulated community by examining the (C, P) values associated with LC1 across samples. Overall functional activity in LC1 scales with the amount of externally-supplied resources (color scale in Fig. 4b): in this simple system, the total amount of supplied resources determines the growth of producers of the cross-fed resource, which in turn determines the growth of consumers.

We then performed the same analysis on the LCs of the experimental data from Estrela *et al*. [15]. Figure 4c shows the corresponding (*C, P*) values for each LC. Biomass constraints under facilitation are typically fulfilled in two regimes: at high functional activity, when all taxa with large component entries are present (triangles in Fig. 4c), or at low activity, when these are absent (inverted triangles). By contrast, samples in which facilitating taxa are present but facilitated taxa are absent fall away from the equilibrium line (squares in Fig. 4), indicating that the facilitation constraint is not fulfilled. Across all LCs, projections of facilitated taxa display larger variability than those of facilitating taxa (horizontal vs. vertical error bars in Fig. 4c). This supports the interpretation (Section 2.4) that fluctuations accumulate downstream from the supplied resources along the cross-feeding chain.

A least component that is not fulfilled reflects an over- or under-representation of taxa relative to the biomass-conservation constraint associated with each resource, which in extreme cases is consistent with extinction of functional—such as consumer—taxa. Within the consumer-resource interpretation, this departure from the constraint is expected to result in in the accumulation of an internally produced resource. Projections onto least components therefore cluster distinct functional states across sampled communities, corresponding to different ways in which biomass constraints are fulfilled or not. Beyond this static classification, projections onto LCs can also track how conservation constraints are progressively established as individual experimental replicates reach equilibrium (Fig. SI 18).

### 2.6 Least Components are Sparse and Dominate the Inferred Coupling Network in Natural Microbial Communities

We next examined LCs in natural microbial communities using several natural datasets drawn from a previous metastudy [23]. Across all datasets, we identified the presence of multiple LCs (purple numbers in Fig. 5a). These components are generally sparser than PCs, involving a smaller effective number of participating taxa in most communities (boxplots in Fig. 5a; Methods). Figure 5b illustrates this sparsity for two representative examples of LCs in the natural soil community shown in Fig. 1g.

**Figure 5.**
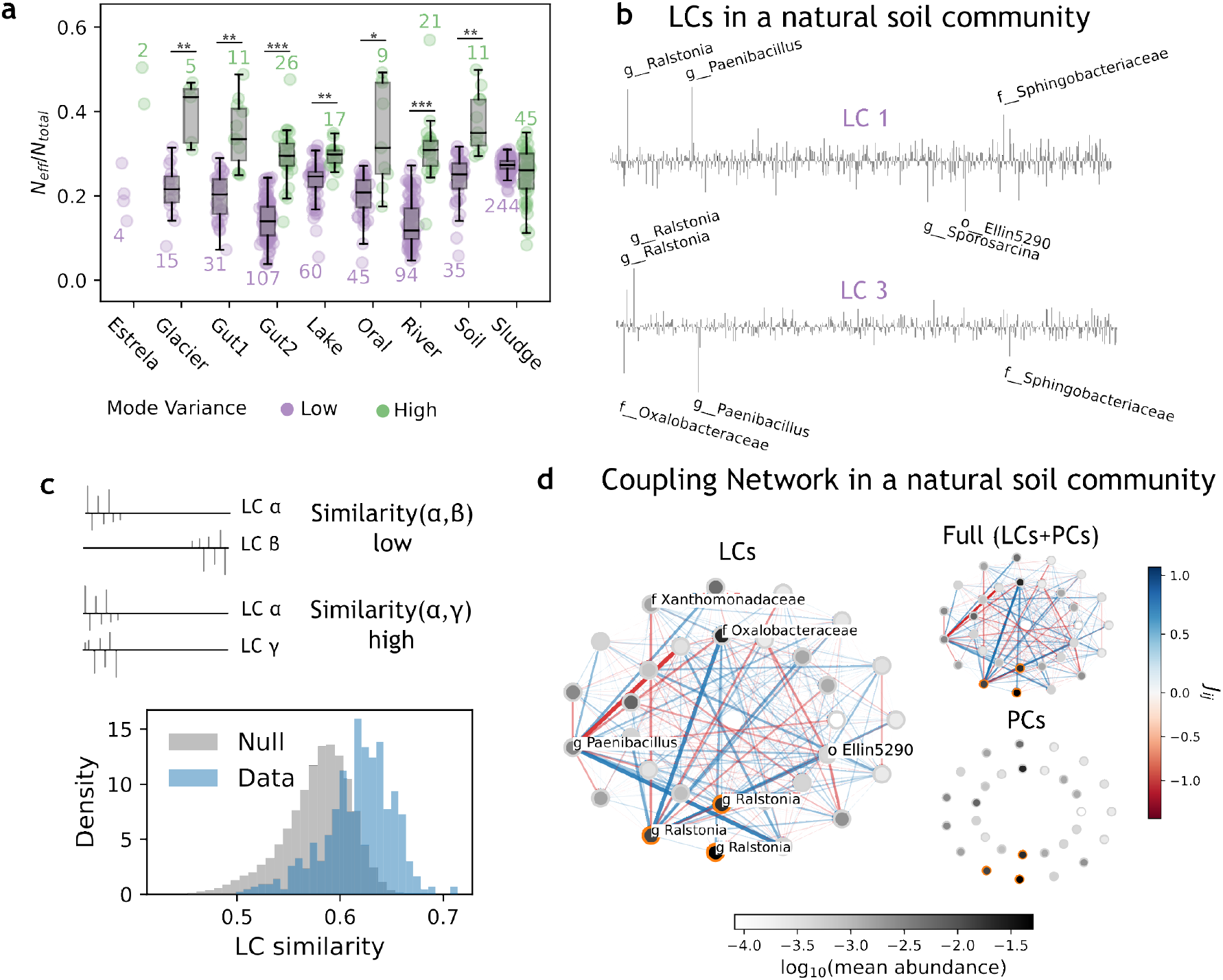
Least Components are sparse and dominate the coupling network of natural microbial communities. **(a)** fraction *N*_*eff*_ /*N*_*total*_ (Methods), of taxa participating in the least (purple) and principal (green) components of multiple datasets of natural microbial communities. Numbers indicate the number of LCs (purple) and PCs (green). The fraction of participating taxa is lower in LCs than PCs across most natural datasets (2-sample Kolmogorov Smirnov (KS) test p-values: < 10^*−*2^ (*), < 10^*−*3^ (**), < 10^*−*10^ (***)). Results for the experimental dataset of Estrela *et al*.[15] are shown for reference. **(b)** example LCs in the natural soil community. The 5 taxa with strongest entries in each LC are labeled. **(c)** Similarity of empirical LCs is larger than expected in the natural soil community (2-sample KS test *p*_*val*_ < 10^*−*130^). **(d)** Coupling network among the 30 taxa that participate in the strongest couplings in the natural soil community. *Ralstonia* taxa (orange edges) and the taxa with which they have strongest couplings are labeled. The network obtained only from LCs (left), is compared to the ones from LCs and PCs (top-right), and from PCs only (bottom-right). All networks are shown in the same color scale.

The presence of LCs with close eigenvalues (Fig. SI 19) suggests that entries on individual LCs are fragile—*i*.*e*. sensitive to small perturbations of the data (Fig. SI 20; Methods). As a result, individual LCs cannot be directly associated to specific ecological interactions. By contrast, LCs in the experimental dataset analyzed in Sec. 2.6 were verified to be robust to perturbations (Fig. SI 20). The fragility of LCs is generally incompatible with a strong sparsity, unless LCs have large entries on overlapping sets of taxa. This condition is satisfied in natural datasets (Fig. 5c for the soil natural dataset, other datasets not shown; Methods), indicating that the LCs consistently involve a restricted subset of taxa.

To bypass the fragility of LCs, we defined an effective coupling network between taxa by averaging contributions across all LCs [22, 25]. The resulting coupling matrix:

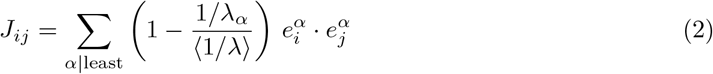

quantifies the extent to which variations in the abundance of taxon *i* are associated with variations in taxon *j*, and viceversa. Here, 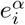denotes the entry of taxon *i* on LC α, and 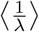 is the mean inverse of nonzero eigenvalues, which weighs each component relative to the null model (Fig. 1). Figure 5d (left) shows the coupling network for the 30 taxa participating in the strongest couplings.

The matrix *J* provides an estimate of the precision matrix of the data (the inverse of the Pearson correlation matrix), consistent with previous efforts to infer coupling networks in microbial communities [28, 17, 29, 30]. Although contributions from LCs and PCs could in principle be included, we found that the strongest inferred couplings are dominated by the LCs, whereas principal components can be ignored (Fig. 5d), a result that also holds in experimental datasets (Fig. SI 22c). Restricting the coupling network to LCs acts as a form of regularization that mitigates overfitting [25].

It is worth noticing that in the experimental dataset [15], the inferred couplings are strongly negative and dominated by LC1, which encodes the relative abundance nature of the data. Upon removal of LC1, the strongest couplings in the network become consistent with the ecological interactions unveiled by LC analysis in previous sections (Fig 3f and Fig. SI 22b).

In the natural soil dataset, the three taxa belonging to the genus *Ralstonia* are among those with the strongest inferred couplings (orange-edged nodes in Fig. 5d). The original study generating this dataset focused on *Ralstonia solanacearum* as the causal agent of wilt disease in tomato crops [24].

Other *Ralstonia* species have also been reported [31, 32] as strong interactors with *R. solanacearum* in microbial communities [33]. The prominence of *Ralstonia* taxa among the strongest couplings highlights the ability of LCs to uncover functionally important taxa in natural microbial communities. As shown in Fig. 5d, taxa within the same genus have different couplings with one another and with other genus, underlining the relevance of interaction stucture at fine taxonomic resolution—a feature that can be lost upon taxonomic coarse-graining [12].

We further examined natural datasets of the gut and found that LC analysis highlights taxa of the genus *Prevoltella*, in particular species *P. copri*, as prominent contributors to the coupling network (SI Section SI 7). Notably, these taxa—and *P. copri* in particular—have been directly linked to host metabolic phenootypes, including hypertension [34, 35]. Together, these results show that LC structure in natural microbial communities is sparse and dominated by a limited set of functionally important taxa.

## 3 Discussion

High dimensional biological data are typically analyzed by focusing on directions of highest variance [20, 21], relegating the remaining dimensions to noise. Here we demonstrated that, in microbial communities, there exist low variance directions that are statistically significant and contain robust and interpretable biological structure. By combining PCA with principled null models, we identified conserved quantities that reflect restrictions imposed by ecological interactions, offering a new way to infer functional organization directly from abundance data.

High variance directions often capture the imprint of environmental variability (Fig. SI 15) [21], partitioning taxa according to environmental preferences (Fig. SI 15 and Fig. SI 17) [36, 37, 21]. Low variance directions, in contrast, reveal constraints arising from ecological interactions—a taxon that grows to high abundance restricts the abundance of taxa that compete for the same resources, while the abundance of a taxon that produces a resource constrains the abundance of consumers that depend on it. As a result, taxon abundances are constrained to vary within hyperplanes, orthogonal to the least components, and the effective dimensionality of the ecological system is reduced (Fig. 1b). When ecological interactions are resource mediated, least components recover producer–consumer structure and reveal cross-feeding relationships that span multiple taxa. We showed that this consumer-resource interpretation of least components accurately explains observed taxon abundances across experimental communities.

In experimental communities, maintaining constant environmental conditions allows these resource mediated constraints to emerge clearly. In natural ecosystems, although individual least components may not map directly to specific mechanisms, they are consistently detected and larger in number than the principal components. Their prevalence in natural communities, where the number of taxa greatly exceeds the number of samples, suggests the existence of mechanisms beyond simple ecological interactions that drive the low-dimensional structure of the community (Section SI 8). Nonetheless, their sparsity highlights influential taxa and reveals internal structure that is not visible from principal components alone.

While we focused on consumer-resource interactions our analysis does not preclude the emergence of low variance directions from other types of interactions—such as toxin secretion or predator-prey interactions. How and to what extenct such interactions give rise to restrictions in the space of species abundances remains an open question for future work.

Importantly, low variance directions transcend pairwise interactions between community members. They can reflect coordinated relationships among many species that emerge from shared metabolic constraints. In simulated communities (Fig. 3c), the first least component captures a cross feeding involving all taxa. In the experimental dataset [15], another least component identifies a four taxon producer–consumer module that links fermentors and respirators (Fig. 3f). These examples show that least components can reveal ecological structure that cannot be inferred from pairwise analyses alone.

Least components also inherit the fundamental limitations of PCA. Because PCA relies on correlations, it cannot infer causal relationships—such as distinguishing which of two cross-feeding taxa acts as a producer or consumer. We resolved this ambiguity by aligning the sign of a taxon contribution to a given least component based on known taxon roles, either from simulation design or experimental literature [15]. PCA is further limited to linear combinations of species abundances, whereas ecological interactions often give rise to complex, potentially non-linear dependencies. This means that PCA may capture only a linear approximation of the curved manifold in which species abundances vary. Furthermore, PCA imposes orthogonality between components although ecological constraints need not be orthogonal. Consequently, individual least components may not map directly onto single mechanisms, even though the space they span remains highly informative about the underlying ecological structure (Section SI 3). Finally, the null model used to test statistical significance of least and principal components assumes that samples are independent. While this assumption is appropriate for cross-sectional data and experimental replicates, it is generally violated in timeseries data, and components—least or principal—might indicate structure stemming from temporal correlations between samples.

The ability to uncover ecological constraints creates new opportunities for engineering biological systems. Because least components describe directions of minimal variability, modifying the abundance of a taxon with a strong contribution in one of these components provides a means to control the abundances of other taxa. Our analysis of data projections onto LCs (Section 2.5) further indicates that deviations from LC constraints can signal the extinction of key taxa. In turn, this could enable predicting the community state at future times (Fig. SI 18). In addition to inferring community dynamics, analysis of LCs could enable the development of new generative strategies for engineering microbial community composition and function, from suppressing undesirable taxa [38, 39] to optimizing production of metabolites of interest [40, 41, 42].

More broadly, our results show that low variance directions can capture interaction mechanisms in high dimensional systems. Although we have focused on microbial communities, we anticipate that the same approach could be applied to different ecological datasets and in data beyond ecology [22], from transcriptomics to socio-economics, to identify conserved relationships and hidden constraints. Viewing least components as structured features rather than noise thus expands the conceptual and practical scope of dimensionality reduction across disciplines.

## 4 Methods

### 4.1 Co-culture experiments

#### 4.1.1 Bacterial isolates, media, and culturing conditions

Both *E. coli* strains derive from the Escherichia coli K-12 MG1655 background. The reference strain (*E*_*K*_) was obtained from an in-house isolate collection maintained at IAME laboratory and is phenotypically resistant to rifampicin. The fluorescently labelled strain (*E*_*F*_) [43] carries a chromosomal YFP–chloramphenicol cassette under the constitutive λ-phage pR promoter inserted at the *intC* locus (intC::pR-YFP-Cm) and was originally obtained from the Quantitative Evolutionary Ecological and Mechanistic Microbiology laboratory at Institut Cochin.

*Acinetobacter guillouiae* was isolated from a natural soil sample collected in Massachusetts, USA, obtained from the Gore Laboratory (MIT), and subsequently screened within our laboratory’s soil isolate collection. This strain was selected based on its strong growth on M9 medium supplemented with citrate (0.2%) and weak growth on M9 medium supplemented with glucose (0.2%) (Fig. SI 12).

Overnight precultures were grown in Nutrient Broth (NB) containing yeast extract (10 g/L, Becton Dickinson) and soytone (10 g/L, Becton Dickinson), adjusted to *pH* = 7. All subsequent experiments were conducted in M9 minimal medium (11.28 g/L; Becton Dickinson) supplemented with *MgSO*_4_ (2 mM), *CaCl*_2_ (0.1 mM), and a 1× trace-metals mixture, adjusted to *pH* = 6.8, and glucose at the indicated concentration. Media were sterilized by filtration through 0.22 µm cellulose-acetate membrane filters (Corning).

Prior to experiments, frozen bacterial stocks were thawed and streaked onto Tryptic Soy Broth (TSB; MP Biomedicals) supplemented with 2.5% agar (Becton Dickinson) plates and incubated at 30°C for 24h. Single colonies were then picked to inoculate 5mL of NB for overnight preculture at 30°C for 18h with shaking at 180rpm. For CFU counting, experimental cultures were diluted in phosphate-buffered saline (PBS; Corning) and plated on TSB supplemented with 2.5% agar. *E*_*K*_ and *A. guillouiae* were distinguished based on colony morphology.

#### 4.1.2 Competition in co-culture

Overnight precultures of *E*_*k*_ and *E*_*F*_ were washed twice in M9 medium supplemented with 0.2% glucose and adjusted to an OD_600_ of 0.45. To initiate experiments, the two OD-adjusted populations were mixed at the target initial percentage (Fig. 2a), diluted to 3 ×the target initial biomass (Fig. 2a), and 75µL of the mixture was inoculated into individual wells of a 96-well flat-bottom plate (Corning) containing 150µL of M9 medium supplemented with 0.2% glucose. Cultures were incubated for 24 h at 30°C in a plate reader (TECAN Infinite M-Plex) with continuous orbital shaking at 182 rpm. OD_600_ and yellow fluorescence (excitation at 490 nm; emission at 527 nm) were measured every 15 min throughout the growth period.

Fluorescence readings were smoothed using a low-pass gaussian filter (Scipy Ndimage Python Package [44]) to remove high-frequency noise prior to the analysis. The abundance of the *E*_*F*_ strain was inferred by mapping fluorescence measurements to OD_600_ using calibration curves obtained from monoculture growth curves (Figure SI 10). The error bars in the OD measurements of *E*_*F*_ include the contributions from the low-pass filtering and the variability in the OD-fluorescence map (Figure SI 10). The abundance of the *E*_*K*_ strain was measured as the difference between total OD_600_ and the inferred OD_600_ of the *E*_*F*_ strain. OD measurement error was neglected, as it was substantially smaller than uncertainty arising from fluorescence-based inference. Analyses were performed at saturation, defined automatically as the knee-point (minimum second derivative) of the OD_600_ growth curves. The experiment included a total of 80 samples: four replicates per co-culture condition and two replicates per monoculture condition. Wells containing air bubbles at the end of incubation in the plate reader were excluded from analysis to avoid interference with OD and fluorescence measurements (n = 3, and 7 wells for monocultures and co-cultures, respectively). Replicates that did not reach saturation within 24 h—specifically, 4 replicates at the lowest initial OD in the 10% *E*_*F*_ condition and 2 *E*_*K*_ monoculture replicates—were excluded from the analysis, consistent with incomplete glucose consumption.

#### 4.1.3 Cross-Feeding in co-culture

Overnight precultures of *E*_*K*_ and *A. guillouiae* were washed twice in M9 medium and adjusted to an OD_600_ of 1.0. To initiate experiments, 5 µL of each culture were mixed at a 1 : 1 ratio and inoculated into individual wells of a 96-well, 1 mL deep-well plate (Eppendorf) containing 290 µL of M9 medium supplemented with glucose at the concentrations indicated in Fig. 2c. Plates were sealed with an adhesive sterile film (AeraSeal) and incubated for 48 h at 30 °C with shaking at 180 rpm.

Following incubation, cultures were plated for CFU enumeration. Colonies were counted across multiple dilution factors when possible, and CFU/mL was estimated using a Poisson-based estimator as described in Ref. [45]. The experiment included a total of 144 samples: 6 replicates per glucose concentration, for 24 glucose concentrations. Pipetting errors from the 96-well liquid-handling device led to a subset of wells (n = 7) exhibiting anomalously low colony counts despite comparable biomass (OD); these wells were excluded from analysis.

### 4.2 Hypergeometric null model for the co-ocurrence of two taxa

We applied this null model to test the significance of the co-ocurrence between the *Pseudomonas* and *Lachnoclostridium* taxa in [15].

Let A and B be two taxa present in, respectively, *K* and *m* samples of a dataset with *M* samples. In a null model of independent taxa, the probability that they would co-occur in exactly *k* samples is

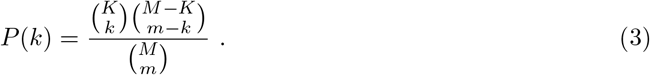

In the *M* = 276 samples of the Estrela dataset[15], *Pseudomonas* is present *m* = 99 times (abundance > 0.01), *Lachnoclostridium* is present *K* = 73 times (abundance > 0.01), and they co-occur exactly *k*^*\**^ = 72 times. The p-value associated to this co-occurence is, under the null hypothesis, the cumulative probability *p* = *Σ*_*k*>*k*^*^_ *P(k)* ≃ 3x10^−16^. We chose the abundance thresholds based on the bimodality of the abundance distribution of the taxa [46]. Results do not change significantly if the abundance threshold is set to 0.

### 4.3 Datasets

The experimental and natural datasets analyzed in this manuscript are all publicly available. The dataset of Estrela *et al*. and Goldford *et al*. were obtained following instructions in Refs. [15, 12]. Natural community datasets were obtained from a metastudy [23, 36]. We considered the taxonomic abundances directly available in the corresponding references, and did not re-process raw sequencing reads. For the dataset of Estrela *et al*., we considered for analysis the 14 taxa with largest mean abundance across all samples. For the dataset of Goldford *et al*., we considered all taxa present at least 10 times in each subdataset—with a single supplied resource—to avoid spurious LCs arising from low-occupancy taxa. For natural communities, we considered all taxa with an occupancy above 50% across samples.

### 4.4 Inference of Principal and Least Components

Let 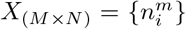 be the matrix of data, where 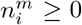 is the abundance of taxon *i* in sample *m, M* is the number of samples (rows) and *N* the total number of taxa (columns) in the dataset. Let 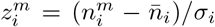 be the z-score corresponding to 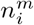, where 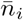 and σ_*i*_ are the mean and standard deviation of the abundance of species *i* across samples, with corresponding matrix *Z*. The empirical correlation matrix of the data is

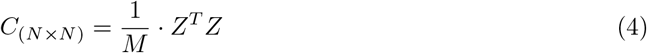

where each entry

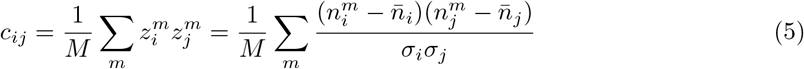

is the correlation between the abundance of species *i* and species *j. C* can be diagonalized as

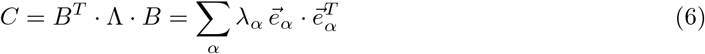

where Λ is a diagonal matrix with entries λ_1_ > λ_2_ > … > λ_+_ > … > λ_*−*_ > … > λ_*N−*1_ > λ_*N*_ ≥ 0 and *B* is the matrix of column eigenvectors, where 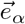 is the eigenvector of eigenvalue λ_*α*_. λ_*α*_ is the variance of the data along the direction 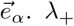 is the smallest eigenvalue of significantly large variance and λ_*−*_ is the largest eigenvalue of significantly small variance, with respect to some null distribution (see below).

Let *r* = *N/M* be the ratio between the number of taxa and samples. If *r* > 1, *C* is rank-deficient and has *N* − *M* + 1 eigenvalues equal to zero, which are not considered. The smallest nontrivial eigenvalue to be considered is thus λ_min(*N,M−*1)_.

### 4.4.1 Theoretical null model: Marchenko-Pastur distribution

If the 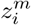 are independent, identically distributed random variables with zero mean and unit variance, the distribution of eigenvalues of the correlation matrix *C* in the double infinite limit *N, M* →∞, with *N*/*M* = *r* fixed, is given by the Marchenko-Pastur (MP) law [47], with bulk density

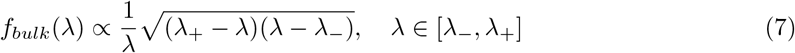

where 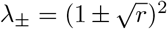 . If *r* < 1, the eigenvalue distribution is fully described by the bulk. If *r* > 1, there is also a Dirac peak at λ = 0 due to the rank-deficiency of *C*.

#### 4.4.2 Numerical Null Model: Basic Shuffling

The simplest numerical null model that we implement consists in shuffling the abundances of each taxon across samples. This is equivalent to sampling abundances from

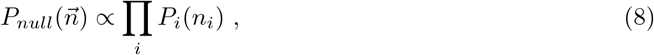

where *P*_*i*_(*n*_*i*_) is the marginal abundance distribution of taxon *i*. By construction, Eq. 8 preserves the marginal distribution of the abundances of each taxon while destroying the correlations between the taxa (see Figure 3b). We use this null model whenever we deal with absolute abundance data, as in the simulations.

This null model approximates the bulk of the Marchenko-Pastur, but has generally wider tails due to finite-size effects. We perform 100 shuffles per dataset, then construct the distribution of the smallest eigenvalue across shuffles and define λ_*−*_ as the bottom 5th-percentile (*p*_*val*_ = 0.05). We define λ_+_ in an equivalent manner from the distribution of the largest eigenvalue across shuffles.

#### 4.4.3 Numerical Null Model: relative-abundance-aware

We now present a null model adequate for datasets of relative abundances. This null model aims at

1. removing the correlations between taxa abundances within samples,
2. preserving the marginal distribution of abundances for each taxon,
3. ensuring that the sum of taxon abundances, *S* =Σ_*i*_ *n*_*i*_, is tightly constrained. More precisely, we assume that the probability distribution of S is given by

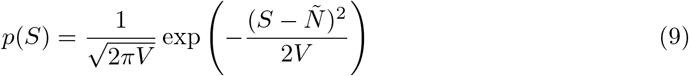

where 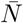 and *V* are the mean and variance of the total sample abundance. In principle, 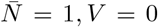, *V* = 0. In practice, since we discard taxa with low occupancy, datasets of relative abundances have 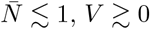, *V* ≳ 0.

Our null model is computationally defined through a Monte Carlo Markov-Chain (MCMC) procedure in the space of surrogate data matrices 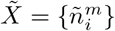, of dimension *M* ×*N*, where the 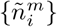 are a shuffled version of the true dataset. Based on (9), we associate to each data matrix an energy

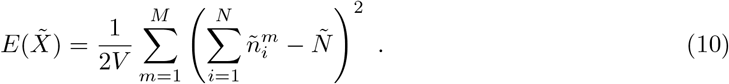

This energy is, up to irrelevant additive constant, minus the log-likelihood of 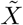 in the null model.

The MCMC procedure is initialized by randomly reshuffling the experimental data matrix *X* across samples for each taxon, hence removing correlations between taxa within samples (requirement 1) and preserving the marginal distribution of abundances (requirement 2). However, the energy *E* has typically very high value with this reshuffled data. To enforce requirement 3, we resort to Simulated Annealing, introducing a fictitious temperature *T*, which is gradually lowered from a high value down to 1.

We run *N*_MCMC rounds_ = 10^6^ rounds. For each round, we randomly pair samples and assign a taxon to each pair of samples, getting a list of proposed swaps. The number of proposed swaps at each step is the minimum between *M*/2 and *N*. We then run through each swap and accept or reject it using the Metropolis acceptance criterion, based on the difference between the energies of the data matrix before and after the proposed swap, respectively, 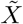and 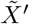:

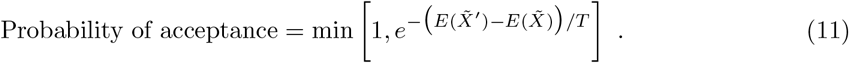

The temperature is initially set to *T* = 50 and progressively lowered to *T* = 1 at *N*_MCMC rounds_/2 and kept to this value for *N*_MCMC rounds_/2 additional rounds. Unless otherwise stated, we generate 100 randomized datasets with this MCMC procedure, to get the same statistical power as in the basic shuffling null model.

Sampling from this complete null model is computationally costly, especially for large datasets, so whenever possible it is preferable to resort to the much simpler basic shuffling. Empirically, we find that this relative-abundance-aware procedure is necessary when *r* < 1 only. When the number of taxa exceeds the number of samples (*r* > 1), as is the case in the natural datasets we analyzed, both numerical null models give compatible eigenvalue distributions (SI Section SI 8), which suggests that the basic shuffling null model can be applied instead.

We define λ_*−*_ and λ_+_ as in the basic shuffling null model, ignoring the least component that arises from the relative abundance structure of the data.

### 4.5 Inverse Participation Ratio as a measure of vector sparsity, and effective fraction of participating taxa per component

The Inverse Participation Ratio (IPR) of a vector, 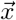, is defined as

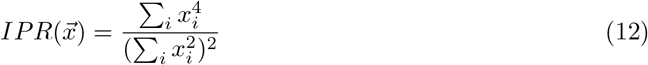

where the *x*_*i*_, *i* ∈{1, …, *N*} are the different vector entries. The IPR ranges from a value 1/*N* for perfectly uniform vectors with all entries equal, to a value 1 for a perfectly localized vector with a single non-zero entry. The IPR can therefore be used to measure the sparsity of the eigenvectors in our datasets, with higher IPR values indicating sparser vectors.

We consider the inverse of the IPR as a measure of the effective fraction of taxa participating in an eigenvector, *N*_*eff*_ /*N*_*total*_ = 1/(*IPR* · *N*_*total*_).

### 4.6 Estimating the fragility of an eigenmode using perturbation theory

If there are two or more eigenvectors of a matrix with the same eigenvalue—*i*.*e*. the eigenvalue is degenerate— any linear combination of them is also an eigenvector of that same eigenvalue. In the context of this manuscript, this implies that least components with degenerate eigenvalue cannot be interpreted as ecological interactions—since they could be arbitrarily combined and remain valid least components. However, the subspace spanned by these eigenvectors remains unchanged.

In practice, since empirical eigenvalues take continuous values, no two eigenvalues will be perfectly degenerate. However, we can assess the fragility of an empirical least component—to which extent its eigenvalue would change if the dataset is perturbed in some way. If the change in the eigenvalue is comparable to the gap with the successive or previous eigenvalue, then we consider that the two modes are effectively degenerate.

Let 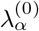 be an unperturbed eigenvalue of the empirical correlation matrix of the dataset, *C*. We consider the perturbation coming from the addition of one new sample, 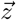, in z-scored abundances. The perturbation to the empirical correlation matrix is

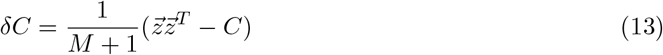

We wish to know what is the expected change in 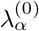, averaged over all possible perturbations 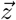. Using well-known results from perturbation theory [48], the first order correction to 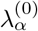 is 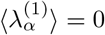, while the second order correction is

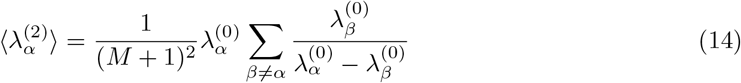

where *M* is the number of samples in the unperturbed dataset. We then compare 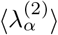 with the difference between 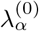 and its nearest neighbor, 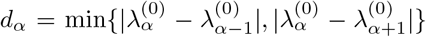. Results for experimental and natural datasets are shown in Fig. SI 20.

### 4.7 LC similarity

We define the similarity of two LCs, *α* and *β*, as

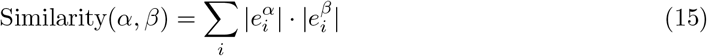

where 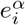 is the entry of the *i*-th taxon on LC *α*. The similarity measure takes values between 0 and 1. For any two LCs, it takes value 0 when all taxa with non-zero entries in one LC have entry 0 in the other. It takes value 1 if all taxa have entries of the same magnitude (up to a difference in sign) in both LCs. A sketch of extreme similarity values is shown in Fig. 5c.

### 4.8 Consumer-Resource Model in batch culture

Let *n*_*i*_ be the abundance of species *i*, and *R*_*α*_ the abundance of resource *α*. Let species and resource abundances change in time according to the equations

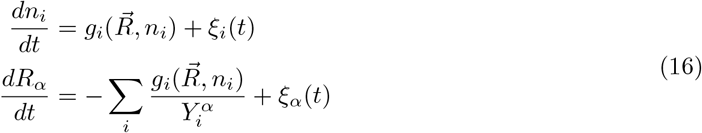

where 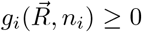 is the growth rate of species 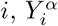 is the yield of species i on resource *α* (> 0 consumption; < 0 production), and ξ_*i*_(t) is white noise with mean 0 and variance *v*_*i*_ (idem for ξ_*α*_, with variance *v*_*α*_). We assume that cell death is negligible. Let us also suppose that the system is supplied with some initial amount of resources and species, {*R*_*α*_(*t*_0_), n_*i*_(*t*_0_) } at time *t*_0_. Inserting the first line into the second in Eq. 16 and integrating until all the resources have been consumed, at time *t*_*f*_, we find back Eq. 1

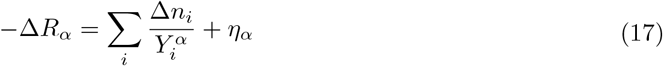

where Δ*R*_*α*_ = *R*_*α*_(t_*f*_) −*R*_*α*_(*t*_0_) ≡ −*R*_*α*_(*t*_0_), Δ*n*_*i*_ = n_*i*_(*t*_*f*_) −*n*_*i*_(*t*_0_) and η_*α*_ is the total contribution of the noise terms in the dynamics (we have omitted the noise term in Eq. 1 for simplicity). Neglecting initial species abundances and rewriting Eq. 17 in vector form, as in Eq. 1, we find that the variance in the projection of the species’ abundances, 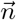, on the vector of inverse yields, 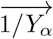, is

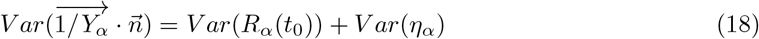

Provided that noise in the dynamics is small enough—*Var*(*η*_*α*_) ≪ 1—and that initial resource abundances are close to constant across samples—*V ar*(*R*_*α*_ (*t*_0_)) ≪ 1—then the variance of 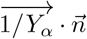 across samples is small and 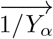 describes a direction of low variance, like a LC.

When inferring directions of low variance from z-scored abundances, every eigenvector 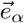 of eigenvalue λ_*α*_ defines an effective vector of inverse yields via

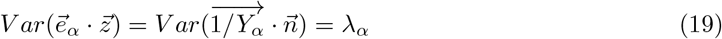

where 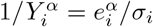. Equations 18 and 19 suggest that the eigenvalue associated to an interaction scales with the fluctuations in the dynamics of the corresponding resource—if fluctuations increase, the restriction on abundances is less tight. We exemplify this in further simulations in batch growth (Fig. SI 11) and in chemostat conditions (Fig. SI 2).

Furthermore, the average change in resource abundance is 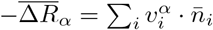.

We performed the simulations shown in the main text using a Monod-type growth function

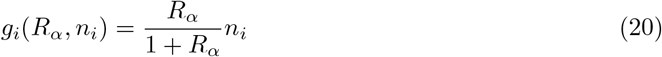

Whenever a resource or a species reach abundance 0 at any point in a simulation, they are kept at 0 (there is no reentry of extinct species or exhausted resources). We used 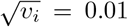 and 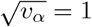.

Initial species abundances were drawn i.i.d from a uniform distribution *U*[0, 1]. The simulated community shown in the main text has 3 resources. Two are supplied from outside the system (resources 1 and 2) and one is produced internally (resource 3). The initial concentrations of the resources in each simulation, s, are given by

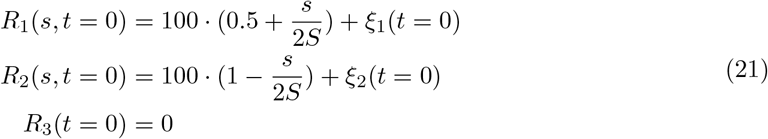

where *S* is the total number of simulations, and ξ_1_, ξ_2_ ∼ *N*(0, 20) are random gaussian variables with mean 0 and standard deviation 20. The choice of initial resource abundances depending on the simulation number s generates diversity in the resources, making some simulations have a larger abundance of resource 1 versus resource 2, while in others the opposite occurs, while keeping initial resource abundances stochastic.

## Data and Code availability

Raw experimental recordings—OD and fluorescence measurements and colony counts—and all Python code used for data analysis and simulations will be made public before publication.

### Acknowledgments

We thank the members of the Amor group for feedback, with special thanks to Matteo Sireci. We are also grateful to Matthieu Barbier for inspiring discussions and Martin Garic for reading the manuscript. This research was funded by the French National Research Agency (ANR) under projects ANR-25-CE45-0796-02, ANR-22-PAMR-0004 (within the France 2030 program), and program ANR-22-CPJ2-0064-01.

## SI 1 Least Components contribute to explaining the structure of high-dimensional data

The inference of PCs and LCs from a high-dimensional dataset can be understood as a problem of maximum-likelihood inference, where the inferred components are those that maximize the likelihood of the observed data. It is possible to calculate explicitly what is the contribution of a single component to this likelihood—i.e. how relevant is the component to explain structure in the data. In this section, we provide a sketch of the derivation of the final results along with some clarifying points. A full derivation of the results can be found in [1].

The components of PCA are vectors in high-dimensional space that point in specific directions, which we call *patterns*. PCs are *attractive* patterns—the data tends to align with them—and LCs are *repulsive* patterns—the data tends to be orthogonal to them. See Fig. 1b for a sketch.

Let 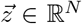 be a sample of z-scored abundances. The probability of such a configuration is given by

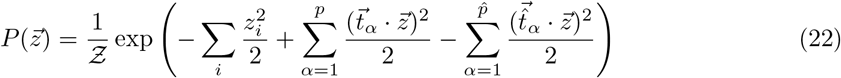

Where Ƶ is known as the partition function and ensures the normalization of *P*. The first term is a gaussian prior on the z-scores of the taxon abundances, while the second and third terms measure the degree to which 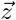 fulfills the different patterns. The second term describes *p attractive* patterns 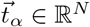, and the third term describes instead 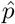 repulsive patterns 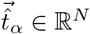. The aim is to infer the set of patterns that maximize the likelihood of a collection of configurations—the empirical dataset. One can safely assume that patterns are orthogonal to each other[1] and that each pattern can be individually paired with a PCA component: attractive patterns with PCs and repulsive patterns with LCs [1].

Each pattern can then be written explicitly from the PCA components as

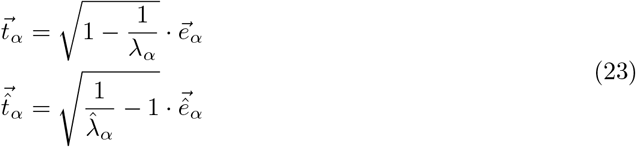

where 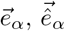 are the PCs and LCs and λ_*α*_, 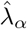 their corresponding eigenvalues [1].

The contribution to the log-likelihood of any one pattern (attractive or repulsive, dropping the hat in 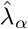) is

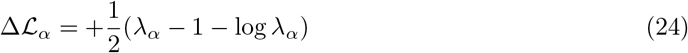

which is strictly non-negative (see Fig. SI 1) [1]. This result in itself showcases the importance of considering modes of low-variance when describing ecological datasets, before their interpretability is considered. In fact, it is perfectly possible that some LCs contribute more to ℒ than some PCs. Table SI 1 shows Δℒ _*α*_ for the empirical eigenvalues of the different PCs in the experimental dataset discussed in the main text [2], showing how the contribution of LCs is comparable, and sometimes larger, than that of PCs. In fact, the biggest contribution to ℒcomes from the component of least variance in the dataset, LC1, corresponding to the compositional nature of the dataset.

**Figure SI 1.**
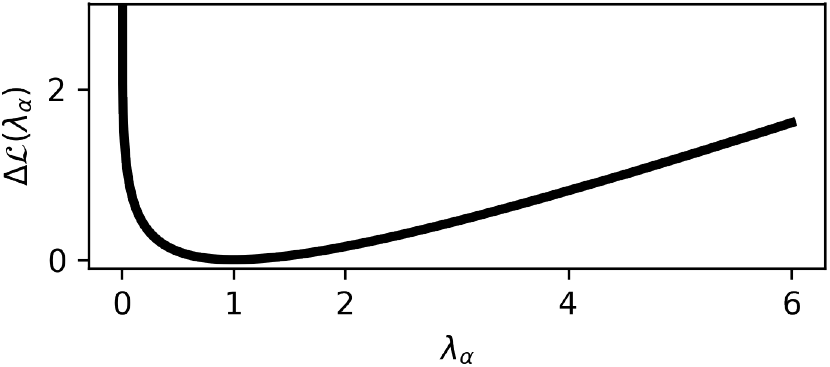
LCs and PCs contribute to the log-likelihood of an empirical dataset. Change in log-likelihood Δℒ as a function of the eigenvalue, λ_*α*_, of a component of PCA. LCs have λ_*α*_ < 1 and PCs have λ_*α*_ > 1.

**Table SI 1.** Contribution of the different PCs to the log-likelihood of the experimental data discussed in the main text [2]. Only eigenvalues incompatible with the Marchenko-Pastur distribution are shown.

| Component ($\alpha$) | $\lambda_\alpha$ | $\Delta\mathcal{L}_\alpha$ |
| --- | --- | --- |
| PC 1 | 3.58 | 0.654 |
| PC 2 | 1.97 | 0.147 |
| LC 4 | 0.489 | 0.102 |
| LC 3 | 0.327 | 0.223 |
| LC 2 | 0.217 | 0.372 |
| LC 1 | $7.40 \times 10^{-5}$ | 4.26 |

## SI 2 Consumer-Resource Model for open systems in chemostat conditions

Here, we describe an alternative version of the Consumer-Resource (CR) formulation for open systems and show that the main results, with respect to the appearance of least components, do not change with respect to batch growth conditions.

Suppose there are *N* species with abundances *n*_*i*_, where *i* denotes species identity. Similarly, there are *R* resources with abundances *R*_*α*_, where *α* denotes resource identity. Species abundances change in time according to

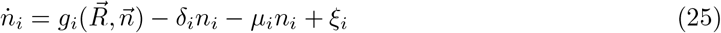

where 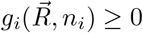 is the growth rate of species *i. δ*_*i*_ is a general dilution (or outflow) rate of species biomass, *µ*_*i*_ a specific death rate and ξ_*i*_ is white noise with ⟨ξ_*i*_⟩ = 0 and ⟨ξ_*i*_ξ_*j*_⟩ = *v*_*i*_*δ*_*ij*_.

The abundances of the resources change in time according to

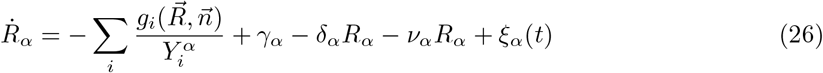

where 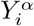 is the yield of species *i* on resource *α* (positive/negative sign indicating consumption/production). *δ*_*α*_ is the outflow rate of resource *α* and *v* its degradation rate, γ_*α*_ its entry rate and ξ_*α*_ noise with ⟨ξ_*α*_⟩ = 0 and ⟨ξ_*α*_ξ_*β*_⟩ = *v*_*α*_δ_*αβ*_.

Assuming the outflow rate is the same for all species and resources, *δ*_*i*_ = *δ*_*α*_ = *δ* ∀*i, α*, and that mass outflow in the open system dominates over death and degradation, *µ*_*i*_, *v*_*α*_≪ *δ*, we can define a generalized biomass for each resource

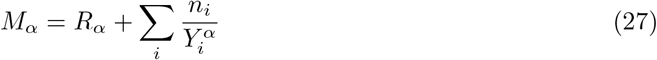

where *M*_*α*_ is a measure of the amount of biomass related to resource *α* present in the system, originating from outside the system, either in the form of the actual resource or in the form of species abundance. Substituting Eq. 25 into Eq. 26, the different *M*_*α*_ follow the stochastic differential equation

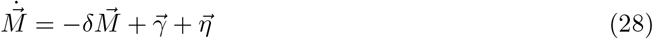

where 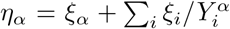, with ⟨*η*_*α*_⟩ = 0 and 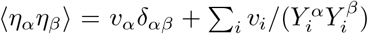. Assuming that *v*_i_ = v ∀*i* and that the vectors of inverse yields of two different resources are orthogonal to each other (see SI Section SI 1), the last expression reduces to 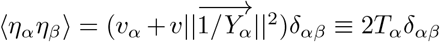.

Then, there exists an equilibrium distribution 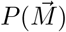

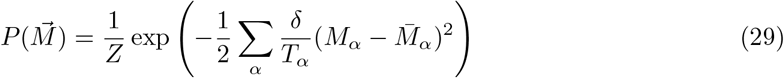

where 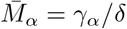.

Equation 29 is a probability distribution for the generalized biomasses *M*_*α*_, and does not define a unique joint probability density on the species and resource abundances. Assuming a gaussian prior on abundance fluctuations, we can write the full joint distribution for the z-scored abundances of the species and the resources as

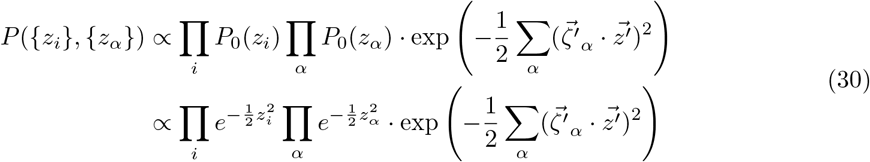

where 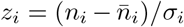 and 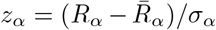 (Latin-alphabet sub-indices indicate species abundances and Greek-alphabet sub-indices indicate resources). 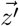 includes both species and resource abundances and we rewrote the generalized biomasses as

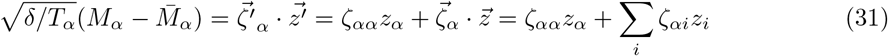

where 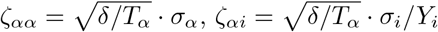 and the vectors 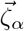and 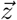 run only over the species’ indices.

Since we only observe species abundances, we can get their joint marginal distribution by integrating out the resources, which leads to

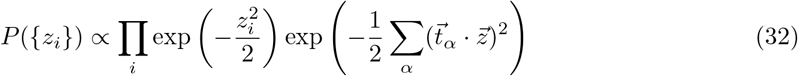

Where 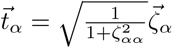

Equation 32 is the same as Eq. 22: in principle, the presence of a resource gives rise to a repulsive pattern, *i*.*e*. a least component, which will be inferrable provided fluctuations are small enough.

We can write explicitly the value of the entry of one pattern in terms of the parameters of the CR model:

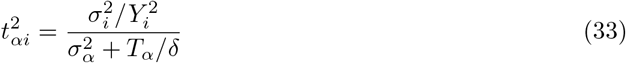

where we recall that 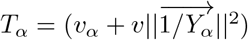. Using Eq. 23 and summing over all the entries of the pattern, we can also predict what is the eigenvalue associated to the least component:

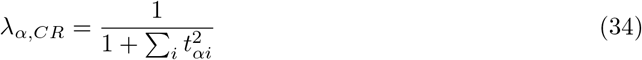

Figure SI 2 shows empirical values of λ_*α*_ and 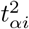, compared to theoretical predictions, for simulations with a single resource and different fluctuations in the resource dynamics, *v*_*α*_. For the other parameters, we used *δ* = 0.1, *v* = 10^*−*6^, *Y*_*i*_ = 1 ∀*i*. σ_*i*_ and σ_*α*_ are measured on the simulations.

**Figure SI 2.**
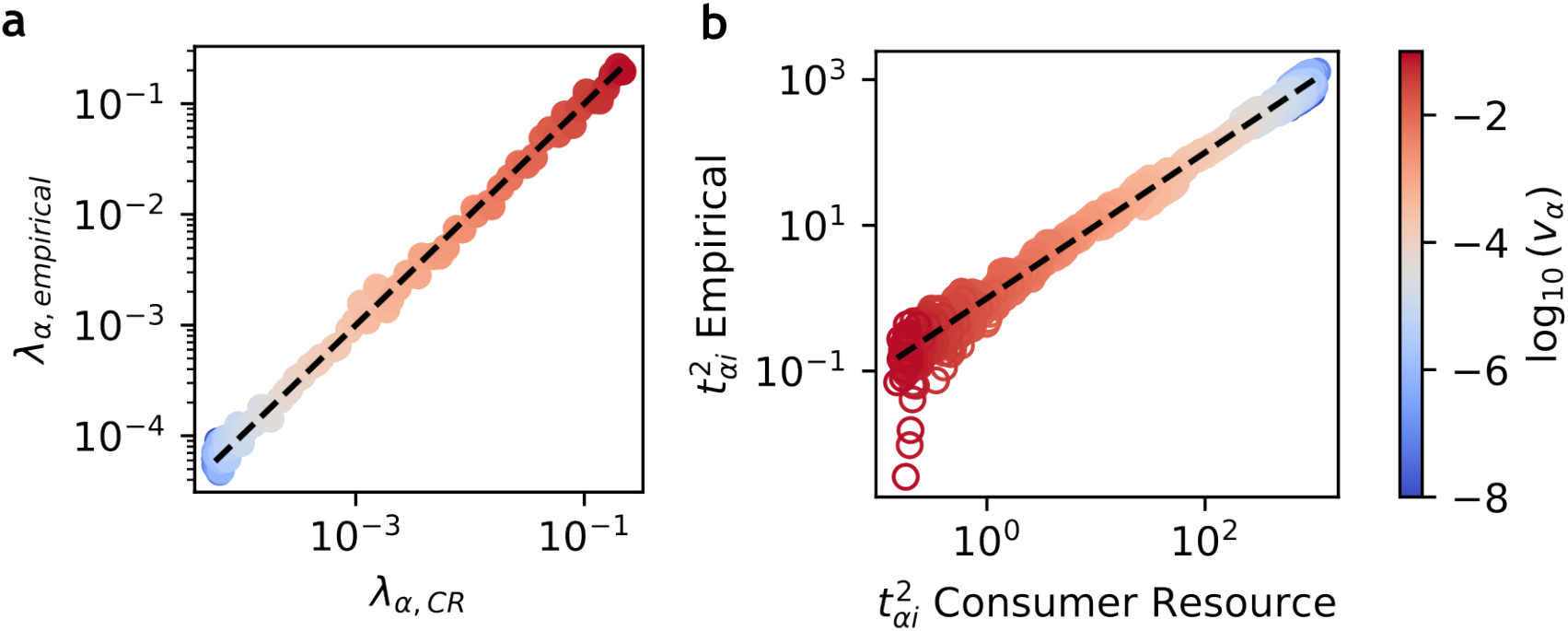
Empirical Least Components are predictable from the parameters of a CR model in chemostat conditions. **(a)** Predicted vs empirical eigenvalue of the least component. Each dot is the smallest eigenvalue in a dataset with *N* = 20 species and *M* = 100 samples. Colors are as in panel B. **(b)** Predicted vs empirical pattern entries. Each dot is one pattern entry in a given dataset (there are *N* pattern entries per dataset). Dot color indicates the variance in the resource entry rate, *v*_*α*_, in the corresponding dataset.

We simulated communities with *N* = 20 species and *M* = 100 samples per value of *v*_*α*_. Empirical eigenvalues are rescaled to account for finite size effects, λ →λ/(1 −*r*), with *r* = *N*/*M* [3]. As expected, the eigenvalue associated to a resource-mediated interaction increases with the variance in the fluctuations of resource.

## SI 3 Chemostat simulations of complex communities

In general, for a complex community with multiple resources being produced and consumed, the inverse yields vector of different resources will not be orthogonal. Therefore, any inferred least component cannot be paired with a single resource. However, the low-variance space spanned by the least components should be highly-informative about the space spanned by the inverse yield vectors, which we can quantify by measuring the overlap between the two spaces (Methods).

To test this, we performed *M* = 500 simulations in a system with *N* = 100 species and *N*_*R*_ = 20 resources. Only 5 of those resources are supplied to the system; they are chosen at random, but are the same across simulations (orange arrows in Figure SI 3a). Their entry rates are sampled from a uniform distribution γ_*α*_ ∼ *U*[0, 10]. Every species consumes one of the resources, *α*, and secretes 2 resources from amongst the next 5, {*α* + 1, …, α + 5}. The full matrix of inverse yields is shown in Figure SI 3a. As with the other simulations, *δ* = 0.1 and ξ_*i*_ = ξ_*α*_ = 1, ∀*i, α*.

**Figure SI 3.**
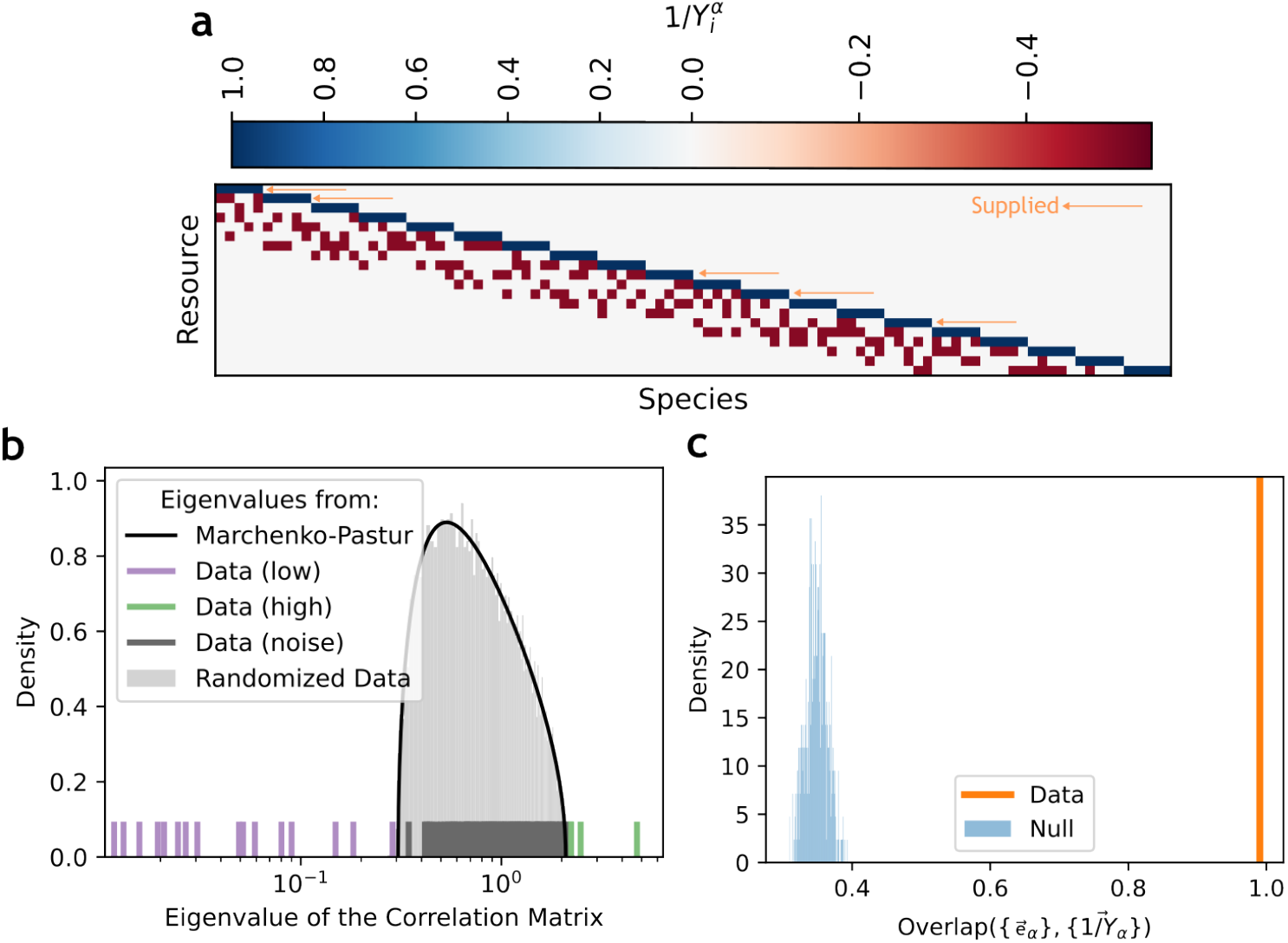
Low-variance eigenmodes recover resource-mediated interactions in complex communities in chemostat conditions. **(a)** Inverse yield matrix of the simulated community. Positive/negative values indicate consumption/production. The 5 orange arrows indicate resources supplied externally to the system in the simulations. The other 15 resources are produced exclusively by the species in the community. **(b)** Eigenvalue distribution of the empirical correlation matrix of the simulated samples. There are 16 eigenvalues falling to the left of the MP bulk: 15 detached and 1 at the edge of the bulk. **(c)** Overlap between the space generated by the eigenvectors, 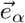, of the 15 detached low-variance eigenmodes, and the 15 inverse-yield vectors,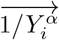, of the resources not supplied to the system.

Figure SI 3b shows the eigenspectrum of the empirical correlation matrix of the resulting dataset. There are 16 eigenvalues of low variance, of which 15 are detached from the bulk and 1 sits at the edge of the bulk. Figure SI 3c shows the overlap of the subspace generated by the 15 detached least components and the 15 vectors of inverse yields for the non-supplied resources. This overlap is close to 1 and significantly higher than the null expectation (shuffling vector entries). This result showcases how the ensemble of least components recovers the resource-mediated interactions in the system, for the cross-fed resources, even when individual least components cannot be paired with a vector of inverse yields.

## SI 4 A taxon with high abundance variance in a relative abundance dataset can give rise to a component of high variance

The null distribution Figure 3b shows a peak of high variance near PC2. We hypothesized that this results from the presence of a taxon with very large abundance fluctuations in a dataset of relative abundances (*Kp* in the dataset), which imposes constraints on the abundances of other taxa in each sample, even in the absence of ecological interactions. In this section, we show how the presence of such a taxon can indeed lead to a PC in a relative abundance dataset.

Consider a hypothetical a dataset with *L* + 1 non-interacting taxa; there are *L* typical taxa and a single atypical taxon which fluctuates significantly in abundance across the dataset. The equation for the relative-abundance-aware null model can be rewritten as

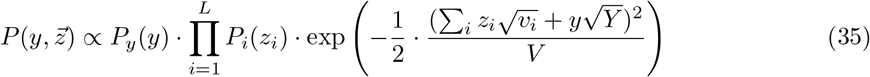

where *y* is the z-scored abundance of the highly fluctuating species,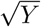 its scale, *z*_*i*_ is the z-scored abundance of one of the typical species and 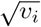 its scale, with *Y* >> *v*_*i*_. *V* is the variance in the total sample abundance: *V* = 0 in theory, but if one neglects rare taxa then *V* ≳ 0. *P*_*i*_ and *P*_*y*_ are gaussian distributions with zero mean and unit variance:

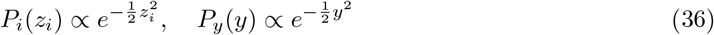

For simplicity, we assume that *v*_*i*_ = *v*, ∀*i*.

We wish to assess for which regime of the parameters *V* and *Y* there will be a LC emerging in the null data and, possibly, a PC due to the highly fluctuating taxon.

The covariance matrix corresponding to Eq. 35 is Σ_(1+*L*,1+*L*)_

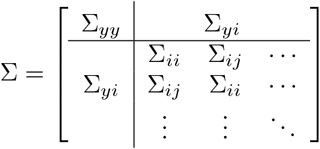

with entries

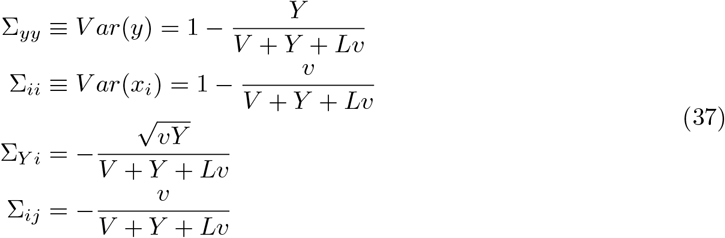

The corresponding correlation matrix of the null data is C_(1+*L*,1+*L*)_

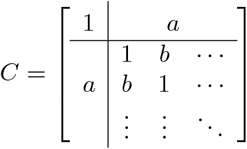

with entries

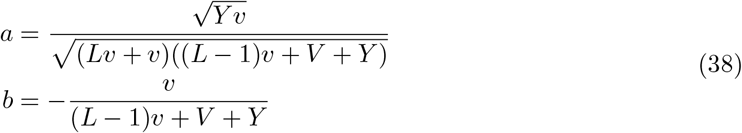

solving the eigenvalue equation for *C* yields two eigenvalues

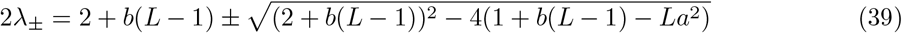

with λ_*−*_ ≤ λ_+_. We can now check the values of *Y* and *V* such that the two eigenvalues cross the significance thresholds, 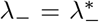 and 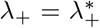 of the null model. This yields two critical curves, *Y*_*±*_(*V*):

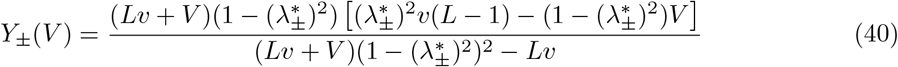

Figure SI 4 shows the (*V, Y*) phase diagram, with all other parameters in Eq. 39 and Eq. 40 set to match those in the dataset by Estrela *et al*.. *Kp* is the highly fluctuating taxon, y. For the other parameters, we took L = 13, and found v from Eq. 37 using the average of the abundance variance of all other taxa, ⟨*V ar*(*x*_*i*_)⟩_*i*_ = 1/*L* ·Σ _*i*_ *V ar*(*x*_*i*_) = 6.11 × 10^*−*3^. In the phase diagram, we show the transition lines given by the edge values of the MP distribution,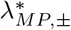, and considering finite size effects, 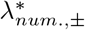. The heatmap shows the average number of significant eigenvalues for different values of *V* and *Y*, across 2000 instances, using 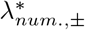 as significance thresholds. Finite size effects consistently shift the significance thresholds towards lower values of V and higher values of Y with respect to the theory.

**Figure SI 4.**
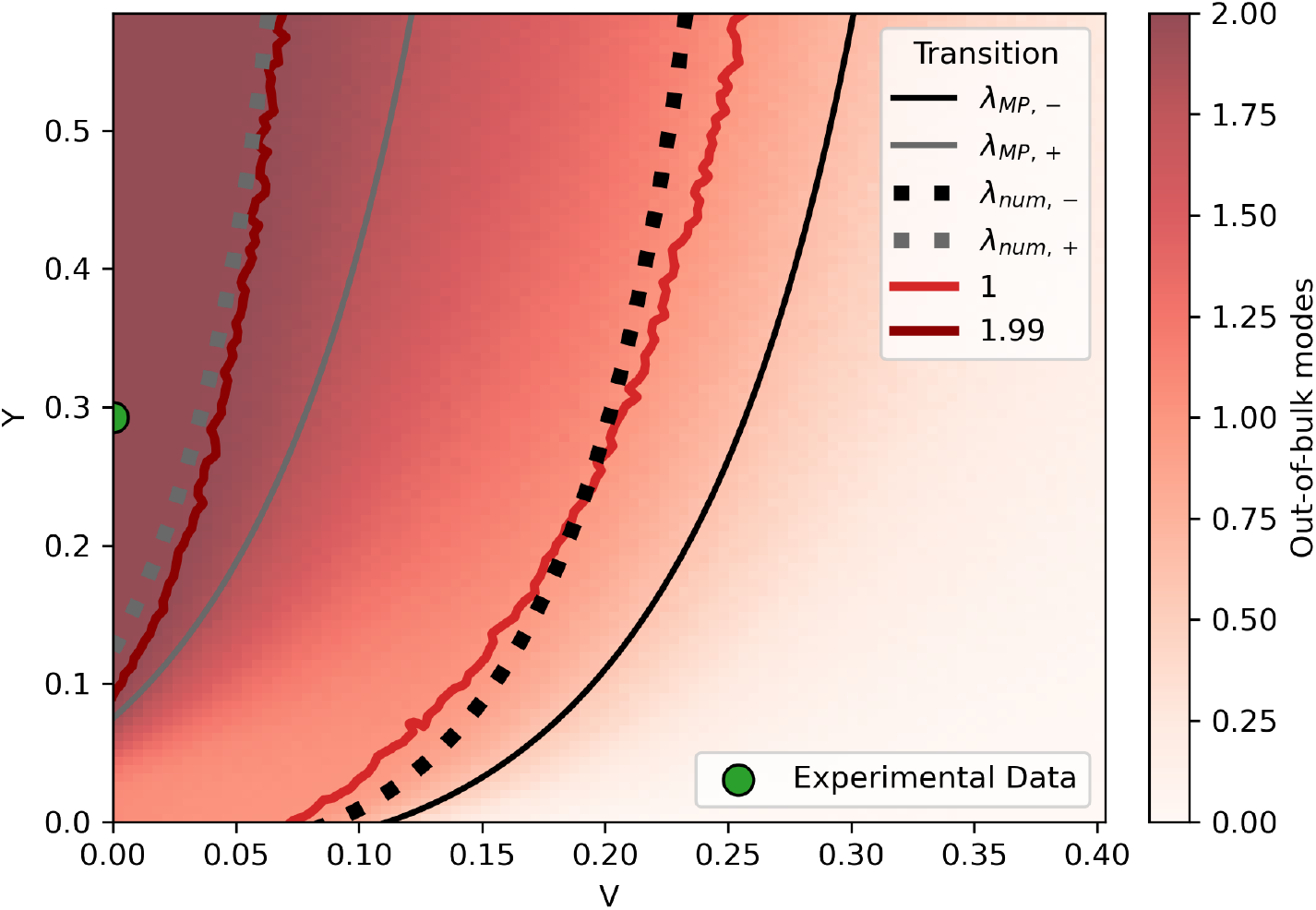
A taxon with high fluctuations in a relative abundance dataset can cause a Principal Component to appear. Phase diagram showing the number of significant high/low- variance modes as a function of the variance in total sample abundance, *V*, and the scale of fluctuations of the highly-fluctuating taxon in the dataset, *Y* . Lines represent the theoretical transition lines when the significance thresholds are taken from the Marchenko-Pastur distribution 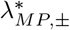 or considering finite size effects 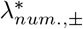. Heatmap shows the average number of significant modes across 2000 simulations, with respect to 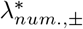. The experimental dataset from Estrela *et al*., analyzed in the main text, falls inside the region with 2 significant modes, which explains the two peaks in the null distribution in Figure 3e.

For decreasing values of V, the 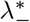 transition always occurs before the 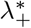 transition. First, there is a LC due to the relative abundance nature of the data (low *V*). Then, if there is a taxon in the dataset with sufficiently large fluctuations (high *Y*), a second transition occurs where a new PC appears.

The experimental dataset of Estrela *et al*. is shown as the green dot in Figure SI 4. We measured *V ar*(*y*) ≡Σ_*yy*_ = 6.33× 10^*−*2^ and obtained *Y* via Eq. 37. The dataset falls deep inside the region where both the low- and high-variance modes are expected.

## SI 5 Inference on other experimental datasets

To test if low-variance eigenvectors can recover ecological restrictions in other experimental communities, we tested our method of Least Components of PCA on another experimental dataset from [4]. In [4], the authors grow microbial communities doing serial growth and dilution cycles, in minimal media supplemented with one of three carbon sources: glucose, citrate or leucine. The authors find that community composition is highly variable at a fine-scale taxonomic level (Amplicon Sequence Variants, ASVs), but more reproducible when coarse-graining composition to the level of Families, for a given supplied resource. Furthermore, family-level composition is predictable from the identity of the supplied resource. As reported in [4], samples supplied with glucose and citrate favor the appearance of taxa in the families *Enterobacteriaceae* (E) and *Pseudomonadaceae* (P), and glucose-supplied samples can be distinguished from citrate-supplied samples by a higher ratio of E to P. Leucine-supplied samples, on the other hand, show a significant amount of *P* but no consistent amount of any other particular family: samples mostly include *Alcaligenaceae, Comamonadaceae, Nocardiaceae, Oxalobacteriaceae* or *Xanthomonadaceae*.

We performed PCA on each dataset, keeping ASVs present in at least 10 samples per dataset. Considering least components, we recovered the results from [4]. Figure SI 5a shows the eigenvalue distribution of the correlation matrix of each dataset against the numerical null and the MP distribution (Methods). Both for citrate- and glucose-supplied samples, there is one eigenmode below the left edge of the MP distribution that captures the stable relationship between *Enterobacteriaceae* and *Pseudomonadaceae* (Fig. SI 5b), grouping most taxa by family on either side of the eigenvector. In the citrate dataset, the eigenmode is statistically significant with respect to the numerical null (*p*_*val*_ < 0.05). In the glucose dataset, it is not significant (*p*_*val*_ = 0.14 > 0.05 against the distribution of left extreme eigenvalues), but we could identify it informed by the results of [4]. In both cases, however, there are multiple significant least components, excluding the LC due to relative abundances: 3 modes in the glucose dataset and 4 in the citrate dataset.

**Figure SI 5.**
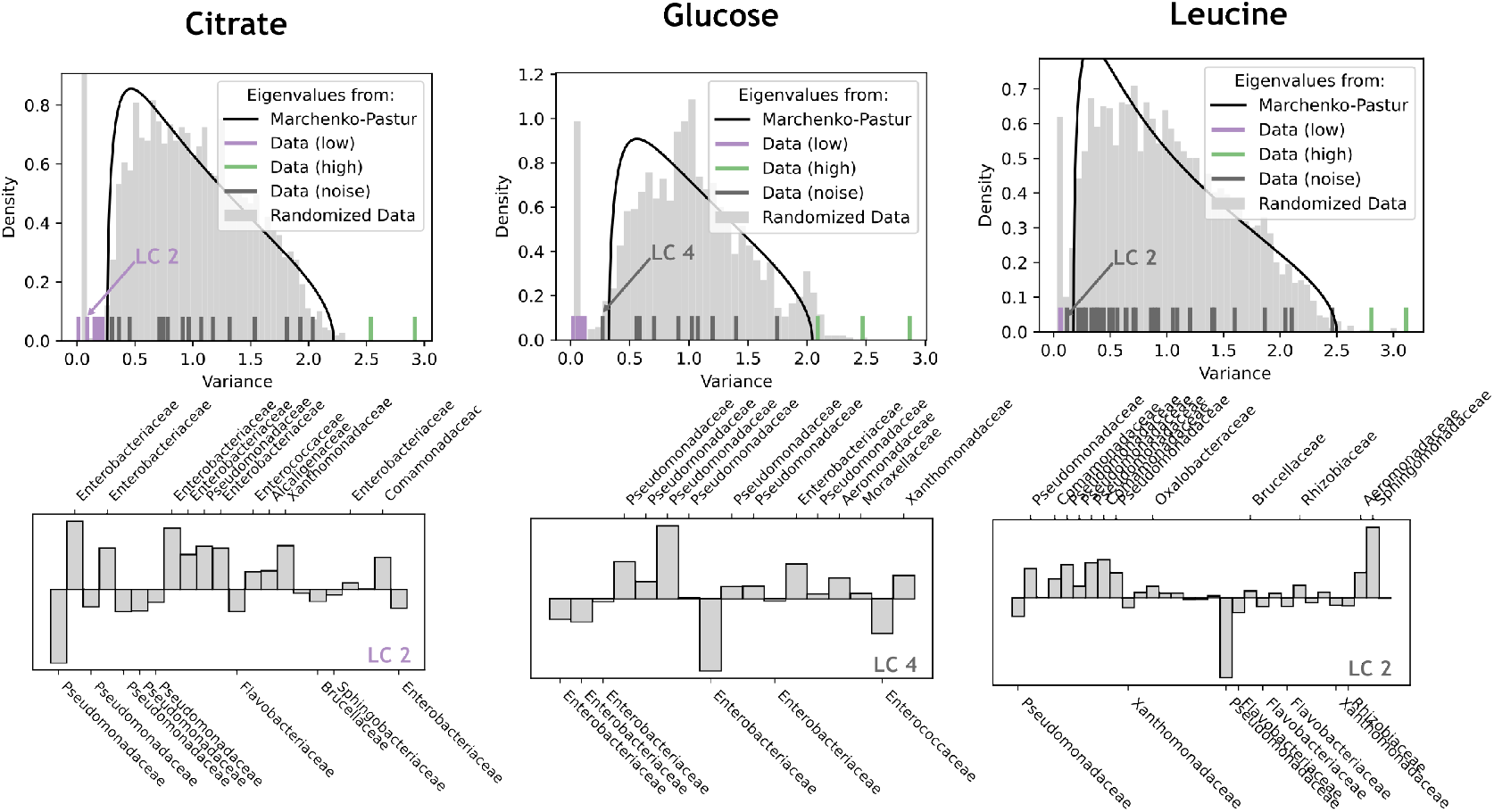
Family-level stable community composition is captured by eigenmodes of low-variance in experimental bacterial communities. **Top:** Eigenvalue distribution of samples supplied with citrate, glucose and leucine, against the corresponding null distributions. **Bottom:** entries on a representative LC per dataset. In the case of the citrate- and glucose-supplied samples, the modes capture the stable composition at the family level reported in [4], placing most *Enterobacteriaceae* on one side and most *Pseudomonadaceae* on the opposite side of the eigenvector (*p*_*val*_ < 0.05 for the citrate component). The LC in glucose sits below the left edge of the MP distribution but is not statistically significant with respect to the numerical null (*p*_*val*_ = 0.14 > 0.05). Communities grown in leucine show no such clear division in [4], and in fact the only significant least component is that for relative abundances (the next component of smallest variance, LC2, is not significant *p*_*val*_ = 0.11 > 0.05, shown).

In the leucine dataset, the only significant least component, with respect to the bulk of the null, is that for the relative abundances. The next LC of lowest variance, LC2, is not statistically significant (*p*_*va*_ = 0.11 > 0.05, shown in Figure SI 5b). This would indicate that family-level composition is not as reproducible in the leucine-supplied dataset as in the other two [4].

A second, high-variance peak, appears in the null distribution of the glucose-supplied dataset. As was the case in the experimental dataset of Estrela *et al*. [2], the glucose dataset considered here contains a single *Pseudomonas* strain whose abundance fluctuations are significantly higher than that of the other taxa (*V ar*(*x*_*P*_) = 0.134 for the *Pseudomonas* strain vs *V ar*(*x*) = 0.053 for the strain with the second highest abundance variance). No such unique taxa exist in the glucose and leucine datasets.

Having considered each of the three datasets separately, we then analyzed them together, by merging them into a single dataset. Only two least components appear (Fig. SI 6a). One corresponds, as expected, to the relative abundances (LC 2, Fig. SI 6b). The other, LC 1, involves only two taxa (Fig. SI 6b) and has lower variance across samples than the relative abundance LC. Both taxa involved appear in most samples at very low abundance, passing the filter on the occupancy, but they coexist at high abundance only once. The cluster at low abundance and the single outlier at high abundance act as two points through which a line can be drawn with very little error, yielding a least component of very low variance that is likely artifactual and has no ecological information. This result highlights the need to properly filter rare taxa from a dataset to prevent the appearance of spurious low-variance modes.

**Figure SI 6.**
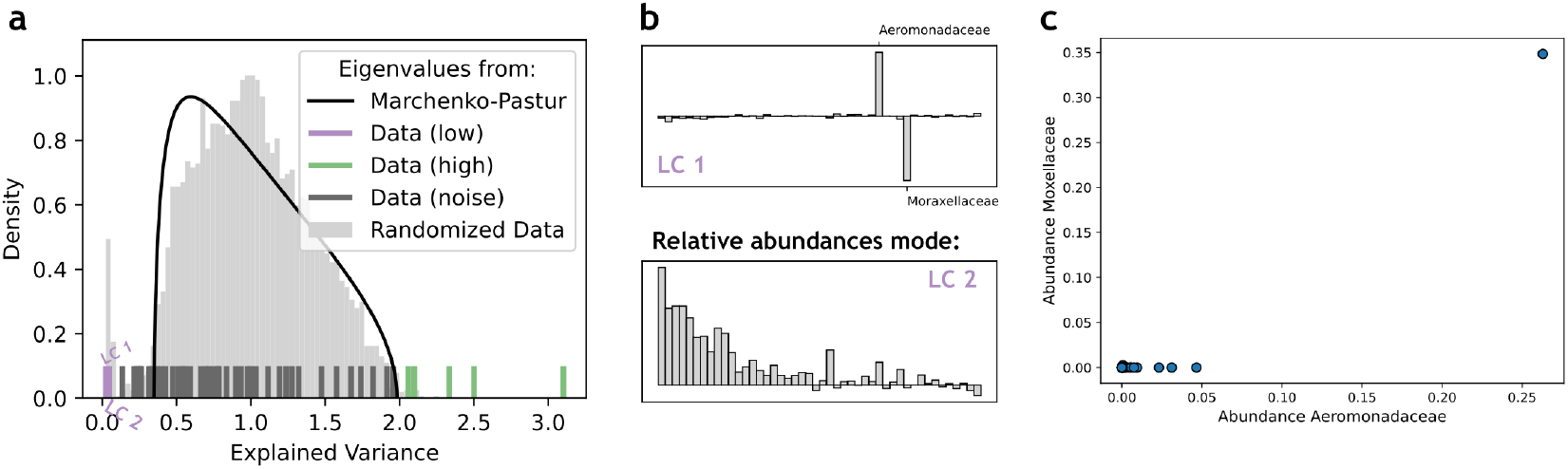
Analysis of the combined datasets of all three resources shows no ecological least components. **(a)** Eigenvalue distribution of the entire dataset. Only 2 modes have statistically significant low variance (*p*_*val*_ < 0.05) with respect to the numerical null. **(b)** The two eigenvectors corresponding to low-variance modes. LC1 (top), the LC of lowest variance, links only 2 taxa. LC2 (bottom) corresponds to the relative abundance LC. **(c)** Abundance of the two taxa in PC45 across samples. Abundances of both taxa is low across most samples, leading to a cluster of points near (0, 0), except for one sample where they both coexist at high abundance. A line can easily be drawn linking the cluster and the outlier, giving a spurious least component.

The analysis of the complete dataset shows no significant LCs with ecological meaning, despite such LCs existed in the citrate and glucose datasets separately. Each least component from those datasets is now fulfilled in just one third of the samples, which corresponds to the supplied resource identity, and is thus no longer constrained enough across *all* samples. To infer interpretable least components, datasets should therefore be collected under identical environmental conditions, which maximizes consistency in microbial interactions across samples.

## SI 6 Change from absolute to relative abundance data destroys some information on least components

To measure how large is the distortion on the data imposed by the change from absolute to relative abundances, we re-analyzed the datasets of natural communities, considering this time the actual sequence counts. Although sequence counts are unreliable measures of true taxon abundance, we can use them as proxy absolute abundances and check how much information is lost in the process of rescaling the dataset to relative abundances. Results show that the number of significant low-variance modes is consistently greater in the case of sequence counts than for the relative abundances (Table SI 2,left). This indicates that some signal is indeed lost in the rescaling to relative abundances.

We then asked whether the signal that remains (the least components inferred from relative abundance data) is informative about the counts data. Since most least components inferred in the relative abundance data represent facilitations (with 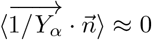 in Eq. 1; Fig. SI 21), and from the discussion that these modes are more robust to the rescaling to relative abundances (see end Section 2.4 and Fig. SI 17), we expected them to be highly informative of the least components inferred from the true counts. To do so, we measured the overlap between the subspaces spanned by the low-variance eigenmodes of the counts and relative abundance datasets.

We define the overlap between any two two subspaces, *A* and *B*, as

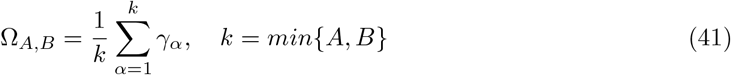

where the *γ*_*α*_ = cos(*θ*_*α*_) are the cosines of the principal angles, *θ*_*α*_, between the two subspaces[5] and can be easily obtained using standard numerical packages [6]. Briefly, the γ_*α*_ are the singular values in the Singular Value Decomposition (SVD) of the matrix 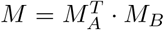 [5], where 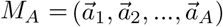 is the matrix of orthonormal column vectors that generate space *A* (idem *M*_*B*_ for space B). Ω_*A,B*_ takes values between 0 and 1. Ω_*A,B*_ = 1 indicates that the two spanned subspaces are equal, and Ω_*A,B*_ = 0 that they are orthogonal.

Overlap across all datasets remained high (Table SI 2,right and Fig. SI 7), indicating that the low-variance eigenmodes in the relative abundance dataset are informative about their counterparts in the true count data.

**Figure SI 7.**
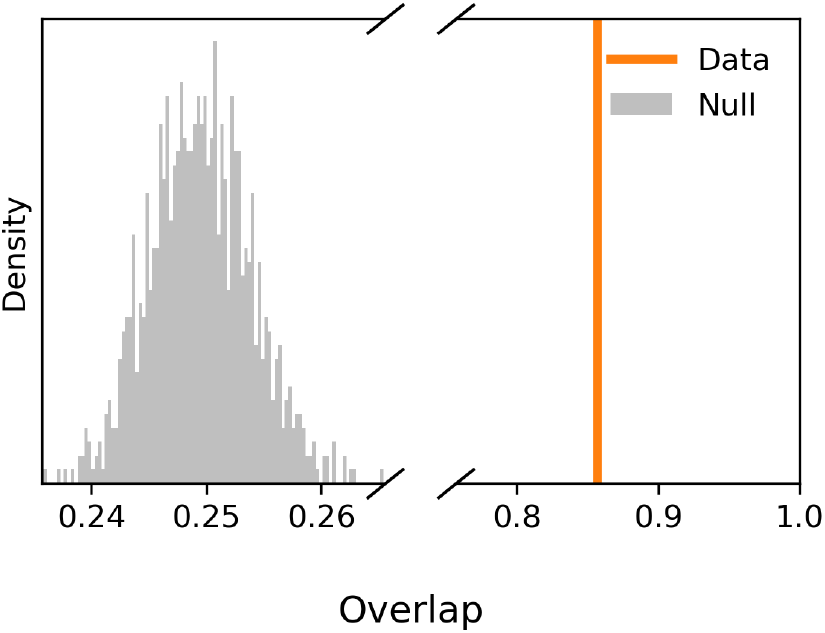
Least components in a relative abundance dataset are informative about the equivalent modes in the absolute abundance dataset. Overlap between the subspaces generated by the least components of the relative and absolute abundance datasets for the soil natural community. The null distribution is obtained by shuffling vector entries in the relative abundance dataset. Summary statistics (true overlap, mean and variance of the null) are as reported in Table SI 2.

**Table SI 2.** Some information is lost by rescaling to relative abundances, although, for natural communities, the information retained is only mildly distorted. **Left:** Across all natural datasets considered, the number of significant modes of low variance is higher when considering the true counts than when considering the relative abundances, indicating to a loss of information when considering relative abundance datasets. **Right:** Overlap between the subspaces generated by the low-variance modes in datasets of relative and absolute abundances, compared against the null overlap when the vectors of one of the subspaces are randomized. Overlaps remain high in all datasets, and significantly higher than null.

| Dataset | Number of Low variance modes |  | Low Variance Eigenspace Overlap |  |
| --- | --- | --- | --- | --- |
|  | True counts | Relative abundances | Data | Null |
| Sludge | 342 | 243 | 0.867 | $0.298 \pm 0.001$ |
| Glacier | 19 | 15 | 0.751 | $0.117 \pm 0.005$ |
| Gut1 | 31 | 31 | 0.843 | $0.149 \pm 0.004$ |
| Gut2 | 113 | 107 | 0.901 | $0.204 \pm 0.001$ |
| Lake | 66 | 60 | 0.928 | $0.240 \pm 0.003$ |
| Oral | 60 | 45 | 0.809 | $0.300 \pm 0.004$ |
| River | 95 | 94 | 0.893 | $0.185 \pm 0.001$ |
| Soil | 44 | 35 | 0.857 | $0.250 \pm 0.004$ |

## SI 7 Analysis of natural gut microbial community datasets

In this section, we provide analyses analogous to the one performed for the natural soil dataset shown in the main text, but for the two gut datasets.

In the manuscript that produced the dataset “Gut1”, the authors present the genus *Bacteroides*, as well as the species *copri* of the genus *Prevotella* [7], as functionally relevant in the gut. In the study that generated the dataset “Gut2”, two enterotypes were identified: one dominated by the genus *Bacteroides* and another by *Prevotella*. The dominance of *Prevotella* was further associated with hypertension [8]. While the authors make no explicit mention of *Prevotella copri* in relation to hypertension, they discuss its role in other diseases [8].

Figure SI 8 shows the coupling networks for the 30 taxa with strongest couplings in each of the two datasets. In both cases, *Bacteroides* taxa appear, in line with the prevalence of this genus in the gut. However, *Prevotella* also appear (nodes with orange edges), in particular for the hypertension study dataset. In this dataset, *Prevotella* taxa represent only 6.9% of the species and 33.7% of the abundance across samples, yet account for 63% of the 30 taxa with strongest couplings in the network. In particular, the *P. copri* taxa are overrepresented, with 8 out of 60 taxa amongst the 30, out of a total of 2003 taxa in the dataset (hypergeometric null model, p_*val*_ < 1.4 × 10^*−*6^).

**Figure SI 8.**
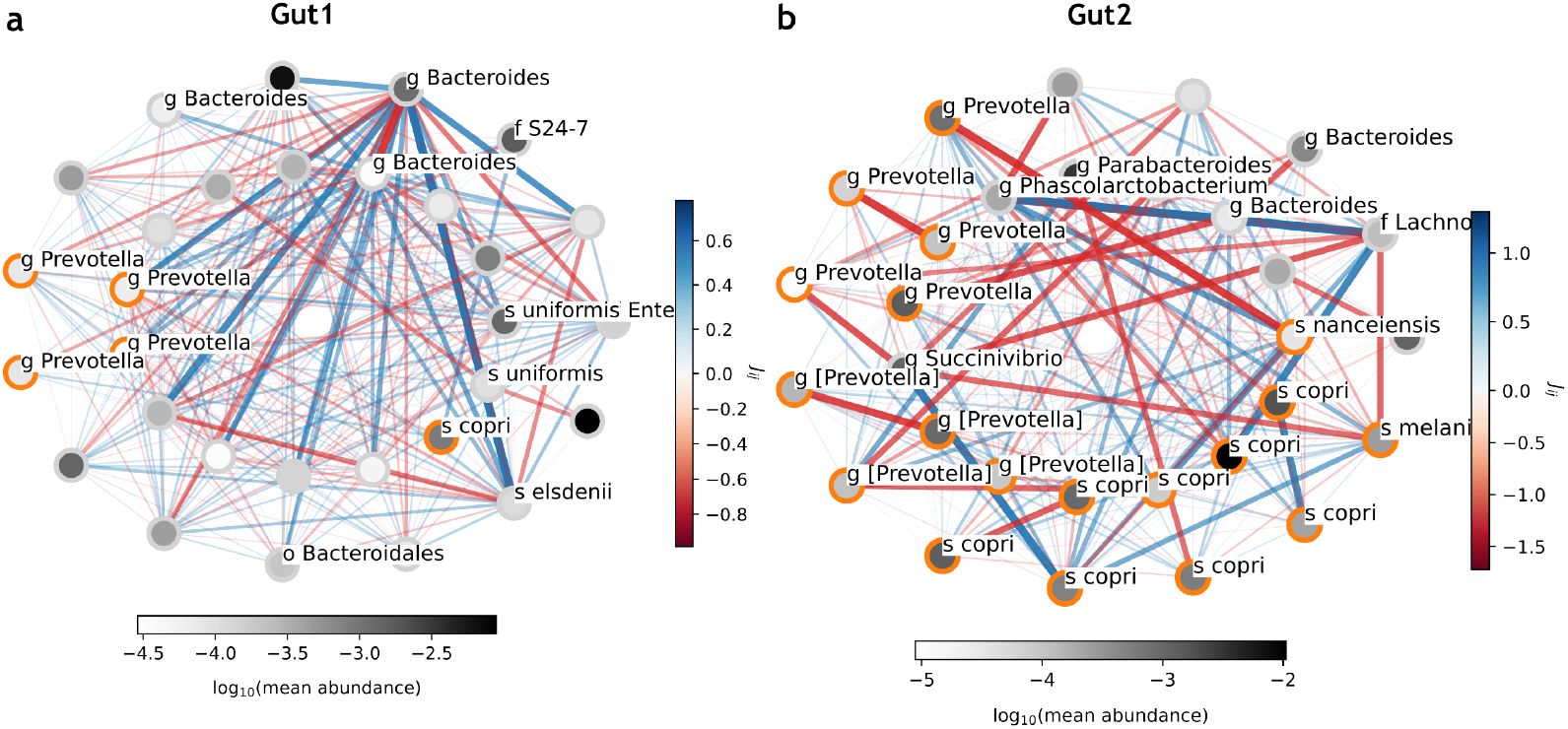
Analysis of the gut datasets. Coupling networks of two gut microbiome natural datasets analyzed in the main text (“gut1” in (a) and “gut2” in (b)). We show the 30 taxa involved in the strongest couplings in the network. Taxa in the genus *Prevotella* have orange-colored edges.

As with the soil microbial community discussed in the main text, the low-variance modes point directly to specific taxa that play a significant role in the community being studied, and here in particular also relate to a diseased state. Furthermore, as was the case with the *Ralstonia* genus in the soil community, the *copri* taxa here do not behave in a coherent way, either with each other or with taxa of other genera. As with the soil community, these results point to the potential usefulness of low-variance modes in highlighting taxa with important roles in natural communities.

## SI 8 Basic shuffling and relative-abundance-aware null model yield the same eigenvalue distribution

Figure SI 9 shows the resulting eigenvalue distribution across 100 runs of the basic shuffling null model and the relative-abundance-aware null model, for the soil natural community. The two distributions are compatible (results are the same for other natural datasets, not shown). This suggests that, with respect to PCA, one can ignore the compositional nature of the data in datasets where the number of taxa exceeds the number of samples. This is consistent with theoretical results that indicate that a component pops out of the null distribution whenever *r* < (λ −1)^2^, where we recall that *r* = *N*/*M* is the ratio between the number of species and samples [3]. When applied to a LC, the smallest possible λ is 0 and the right hand side of the inequality is at most 1, suggesting that no LC should appear in a dataset with more species than samples—*r* > 1. In particular, the mode that encodes relative abundances does not appear in the null model, suggesting that the relative-abundance nature of datasets with *r* > 1 is irrelevant for the inference of LCs.

**Figure SI 9.**
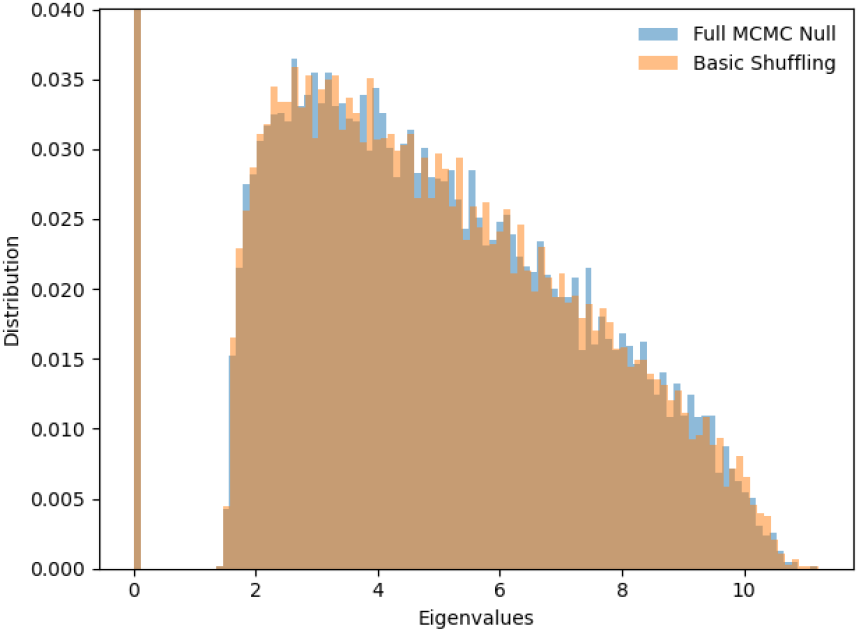
Basic shuffling and relative-abundance-aware null model give the same eigenvalue distribution for datasets with more taxa than samples. Distribution of eigen-values of the randomized Soil natural community dataset, obtained via the basic shuffling null model (orange) and the relative-abundance-aware null model, which accounts for constraints on the sum of taxon abundances in each sample (blue). The two distributions are compatible (2-sample KS test statistic: 0.0036, p-val = 0.874).

The existence of multiple LCs in the natural datasets we analyzed suggests the existence of non-trivial structure in the data. For example, there might be correlations between different samples stemming from a non-trivial spatial organization of the samples. In the soil natural community [9] this might be due to the spatial positioning of plants in the field, and in the gut datasets it might reflect differences in diet between different cohorts [7, 8].

## SI 9 Supplementary Figures

**Figure SI 10.**
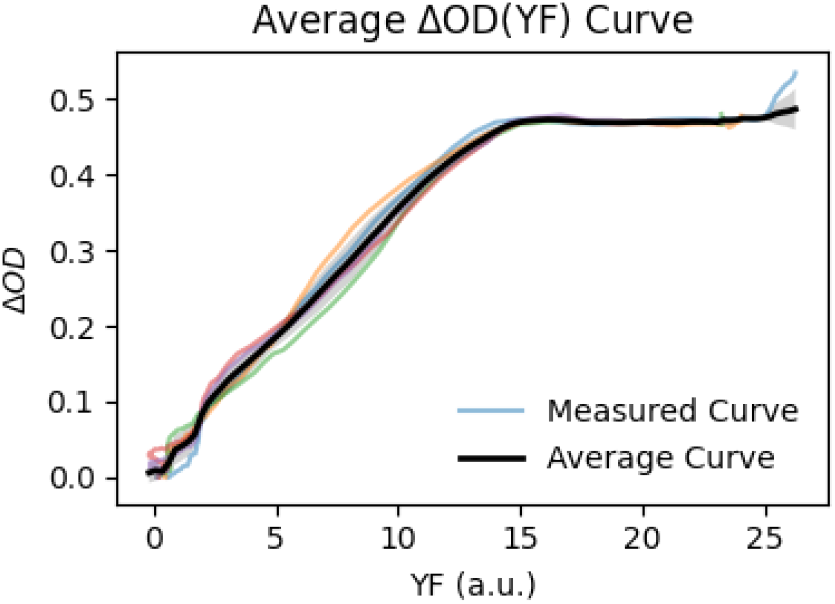
Fluorescence predicts OD below saturation values. OD measured as a function of Yellow FLuroescence (YF) of the *YFP* strain in monoculture. Colored lines are individual measurements. The black line is the average, which we used to predict OD of the *E*_*F*_ strain from YF measurements in the co-culture experiments. Note that, during exponential growth, YF and OD are approximately linearly proportional. After saturation, however, OD flattens but the YF measurement continues to increase, presumably due to additional expression and YFP maturation.

**Figure SI 11.**
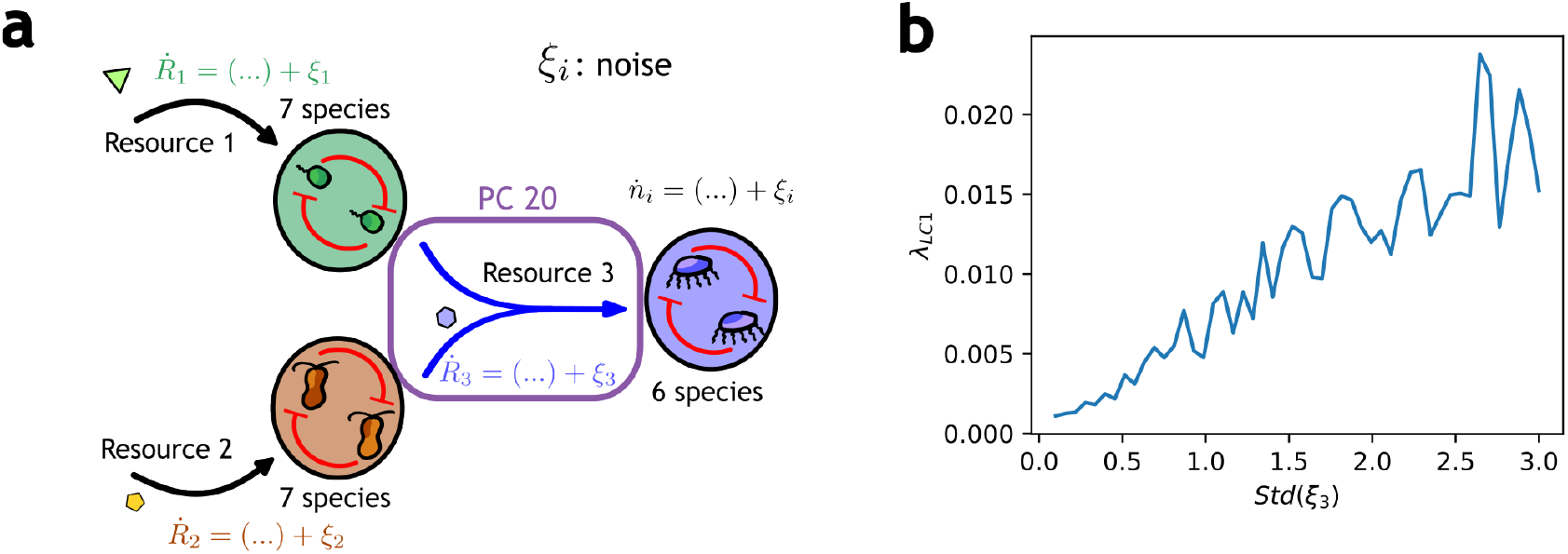
Variance of LCs scales with the fluctuations in the dynamics of the corresponding resource. **(a)** Community structure in the simulated communities is the same as in the main text. We keep the variance of all noises constant across simulations except for ξ_3_. For every value of ξ_3_, we performed 100 simulations. 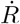 indicates the derivative in time.**(b)** eigenvalue of LC1, plotted against the standard deviation of the noise in the dynamics of resource 3, *Std*(ξ_3_)

**Figure SI 12.**
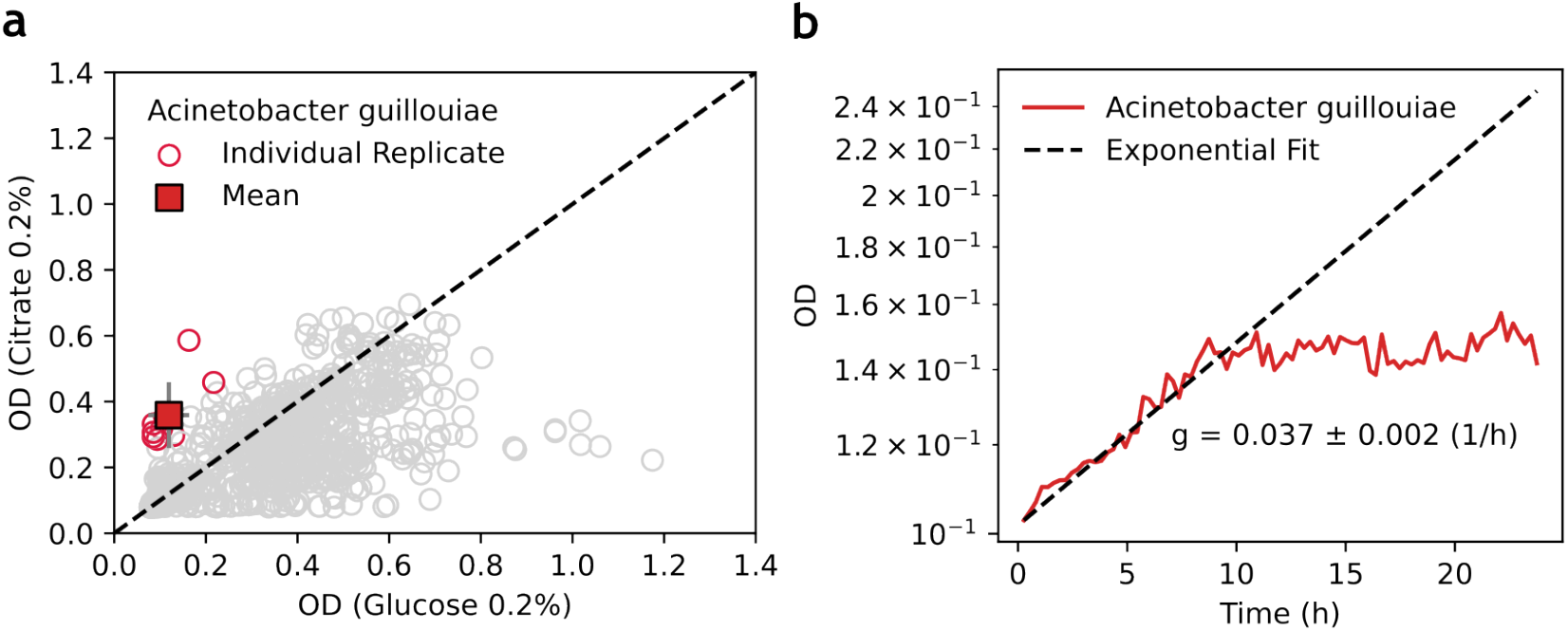
*Acinetobacter guillouiae* grows well on citrate, but poorly on glucose, in monoculture. **(a)** OD measurements at growth saturation of monocultures of all isolates in our bacterial library, grown on either glucose or citrate. Individual dots are individual replicates. Red dots are for replicates of *Ac. guillouiae*; gray dots are for other strains. The square marker is the average of all replicates of *Ac. guillouiae*. **(b)** OD growth curve of *Ac. guillouiae* in minimal medium supplemented with glucose. The dashed line shows the fit to the exponential growth phase, which yields a value of the growth rate of *g* = 0.0037 ± 0.002 h^*−*1^ is the fitted growth rate.

**Figure SI 13.**
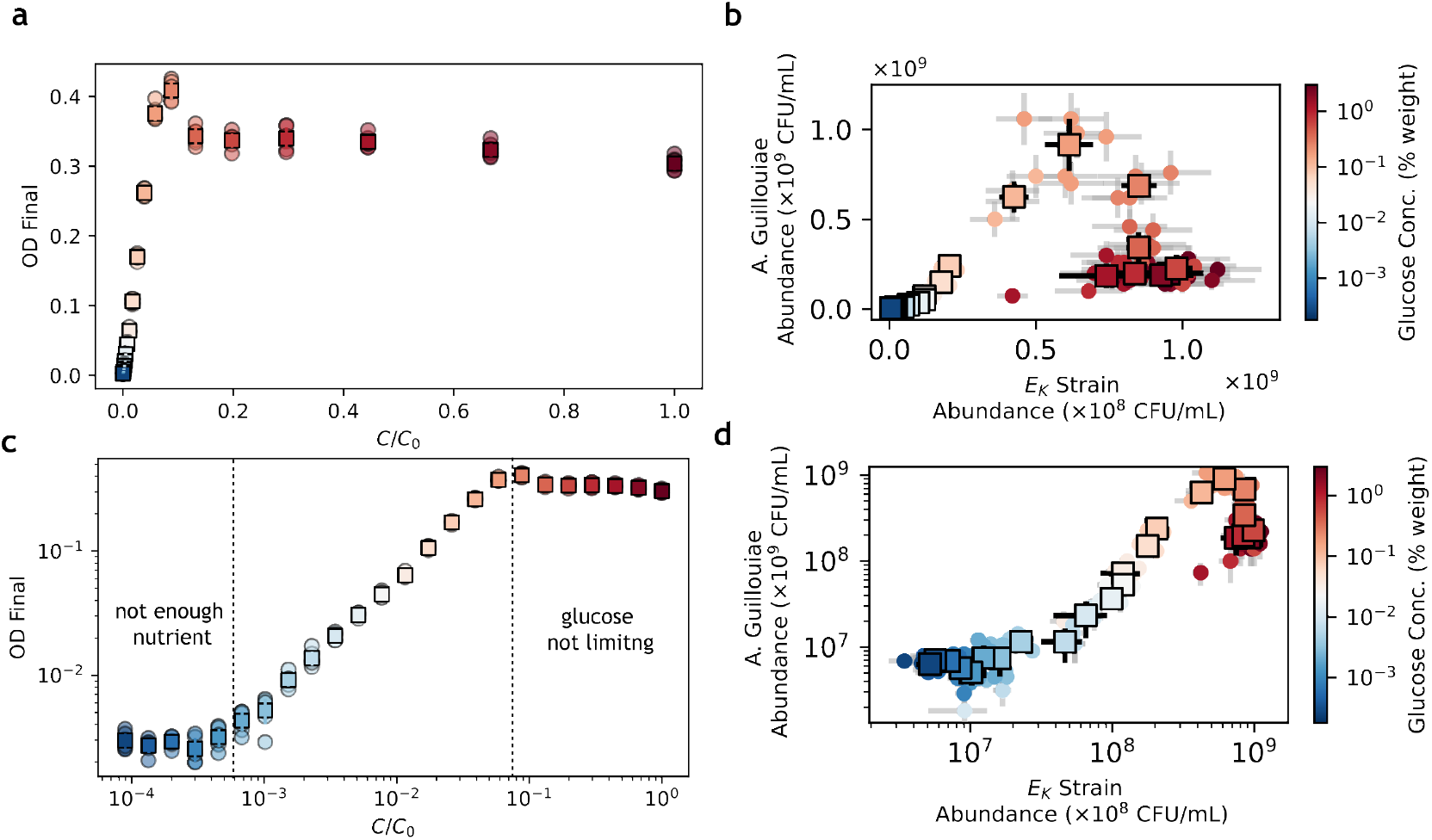
Full results of the two-strain cross-feeding experiment. (**a)** Final OD measurements as a function of initial glucose concentration (expressed as a fraction of the maximum glucose concentration, *C*_0_ = 3% by weight). **(b)** Abundance of *A. guillouiae* versus abundance of *E*_*K*_ in all experimental conditions. **(c**,**d)** same as A,B in log-log scale. **All panels:** dots are individual experimental replicates. Squares are averages over experimental replicates with the same glucose concentration.

**Figure SI 14.**
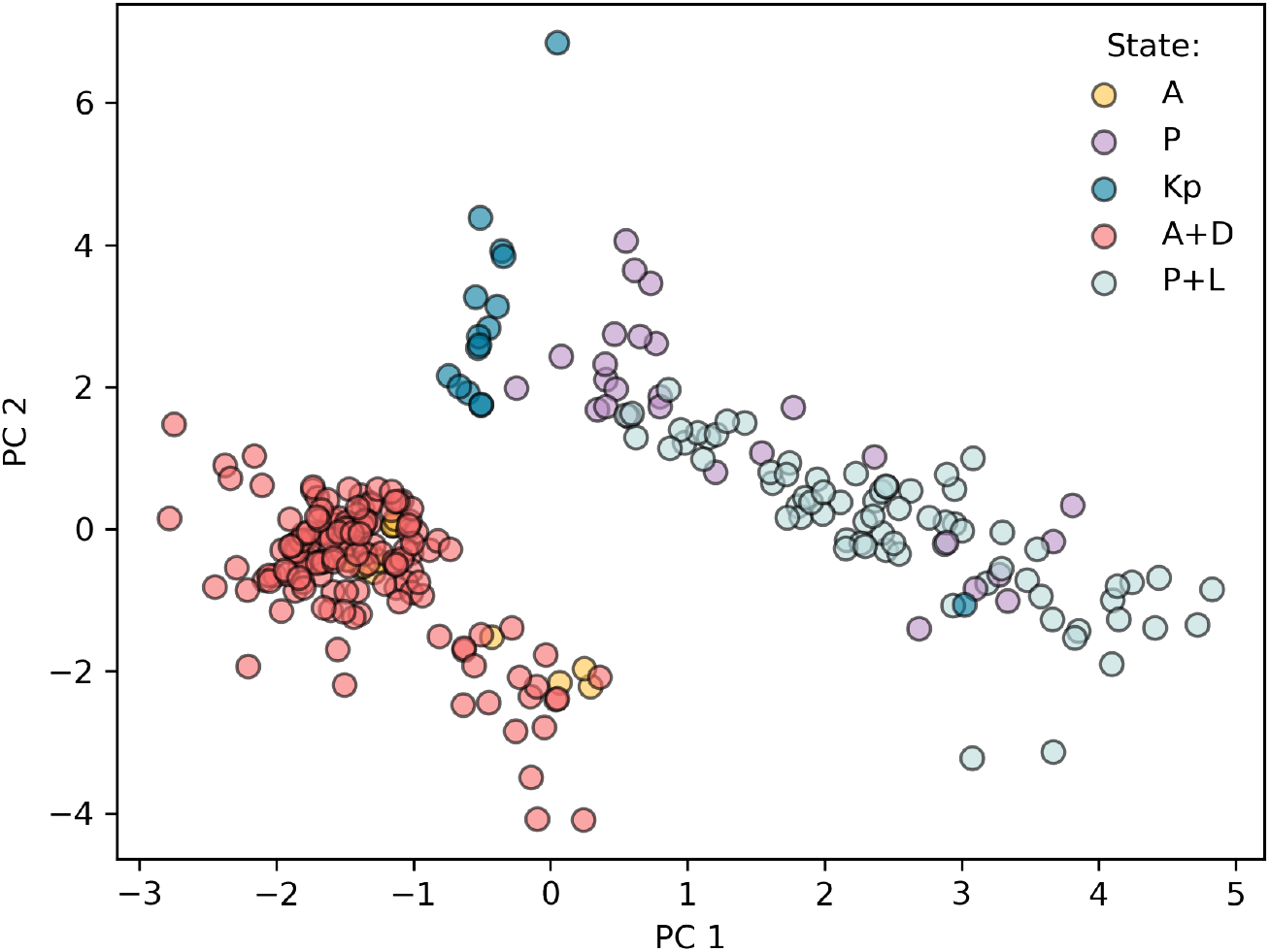
First two principal components reveal the identity of the dominating respirator taxon in the Estrela *et al*. dataset [2]. Data samples projected on the first two principal components of the dataset. Samples are colored and labeled according to the dominating strain at the lowest level in the interaction network in Figure 3b (*A+D* if *Alcaligenes* is the dominating respirator and *Delftia* is present, *A* if *Alcaligenes* dominates but *Delftia* is absent, idem for *P* and *P+L*, and *Kp* if both *Alcaligenes* and *Pseudomonas* are absent in the final communities).

**Figure SI 15.**
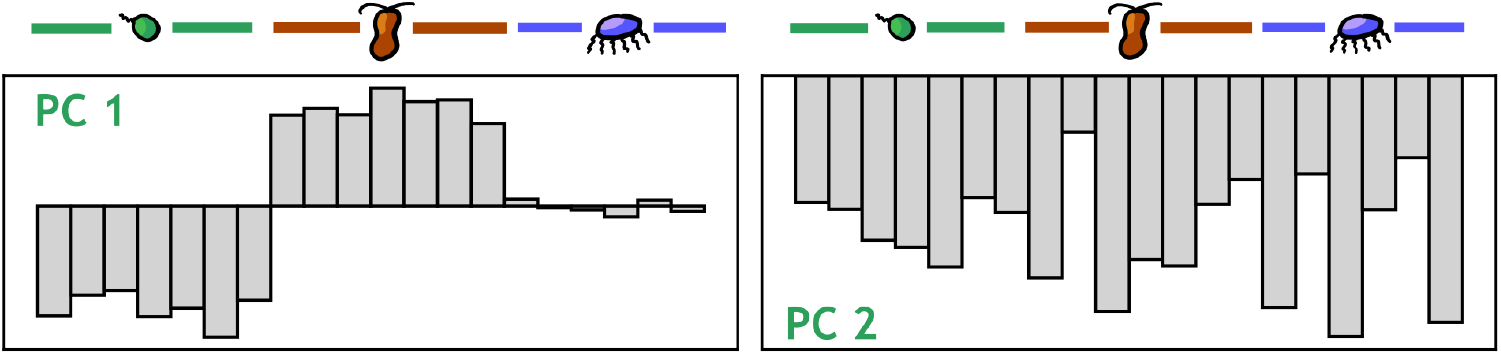
First two principal components for CR simulations in batch growth. **Left:** component entries of individual species on PC1. **Right:** idem on PC2.

**Figure SI 16.**
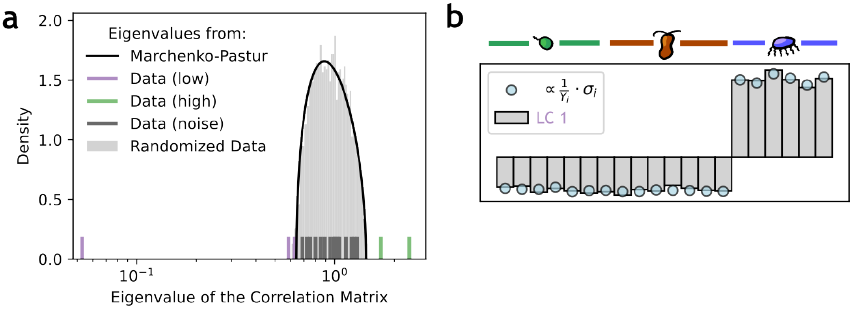
Low-variance modes infer relative inverse yields in chemostat Consumer-Resource (CR) models. **(a)** Eigenvalues of the correlation matrix of a dataset with *M* = 500 samples and *N* = 20 species. Interaction network is the same as in Figure 3. We used *δ* = 0.1, γ_3_ = 0, *γ*_1,2_ ∼*U*[0, 10] and *v*_*i*_ = *v*_*α*_ = 1 ∀*i, α* (see SI Section SI 2 for details). **(b)** Species entries on LC1 (gray bars) and inverse yields on the cross-fed resource, rescaled to units of z-scored abundances (blue dots). LC 1 captures the cross-feeding relation amongst all species via resource 3, as in Figure 3.

**Figure SI 17.**
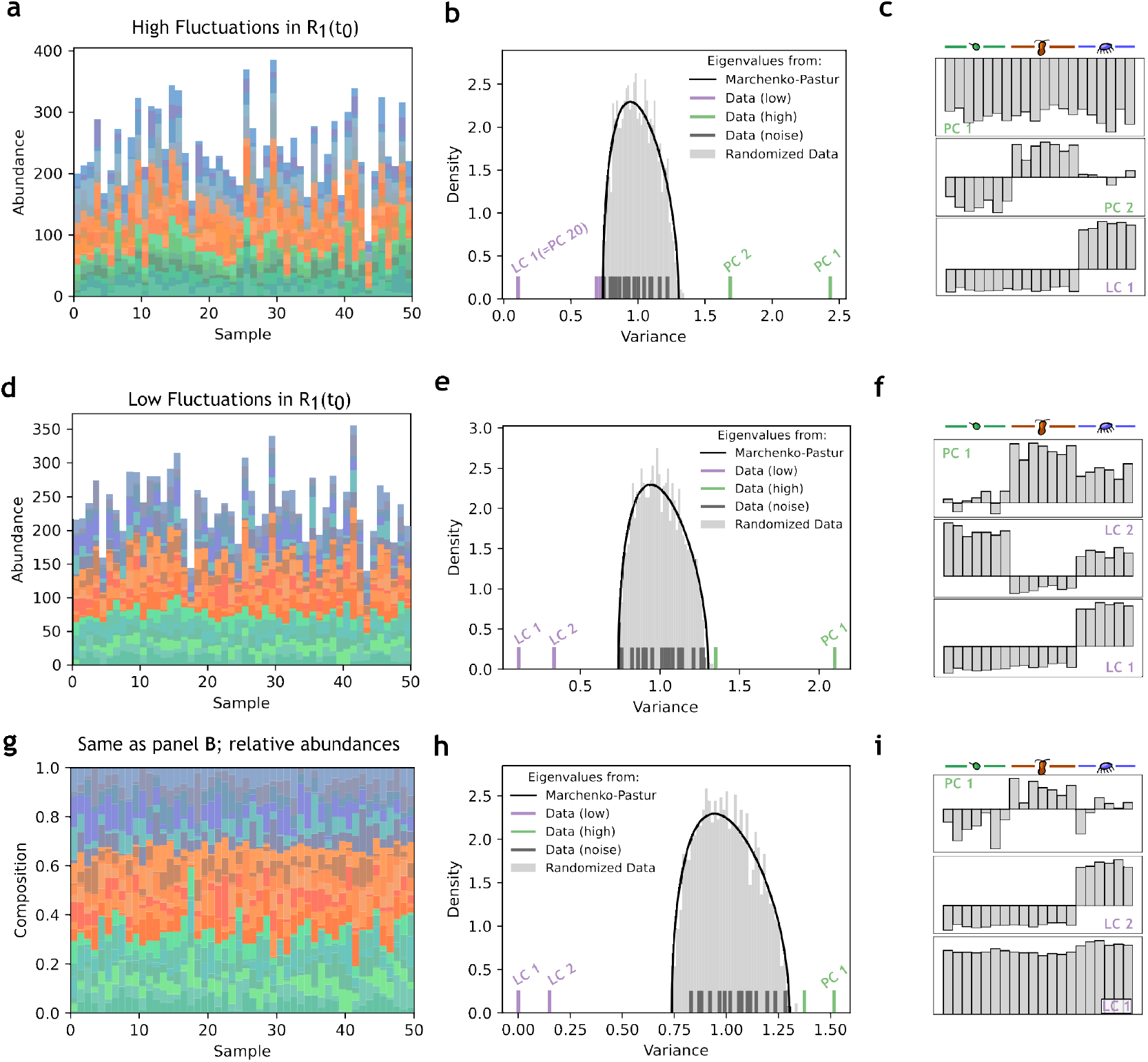
Competitive interactions require small fluctuations to emerge as a LC, and are distorted upon rescaling to relative abundances. **All panels:** the community has the same metabolic structure as in Fig. 3a. Species belong to one of three metabolic groups—green, brown, blue—depending on the consumed resource—1, 2, 3. Resources 1 and 2 are supplied and resource 3 is cross-fed.**(a)** Representative samples in a system with high fluctuations in resource 1 (green) and 2 (blue). **(b)** Eigenvalue distribution of the system. The system has two PCs and one LC. **(c)** LC1 captures the cross-feeding via resource 3. PC1 and PC2 reflect the fluctuations in the two supplied resources, 1 and 2. **(d)** Representative samples in a system with low fluctuations in the entry rate of resource 1 (green). **(e)** same as B, for the system shown in panel D. **(f)** PC1 reflects fluctuations in the influx of resource 1. LC1 captures cross-feeding via resource 3, and LC2 unveils competition for resource 1. **(g)** Same data as D, but rescaled to relative abundances. **(h)** Same as B,E. **(i)** PC1 captures the change in composition depending on the abundance of resource 1 vs resource 2. LC2 recovers cross-feeding via resource 3. LC1 reflects the restriction due to relative abundances. No LC reflects the competition for resource 1.

**Figure SI 18.**
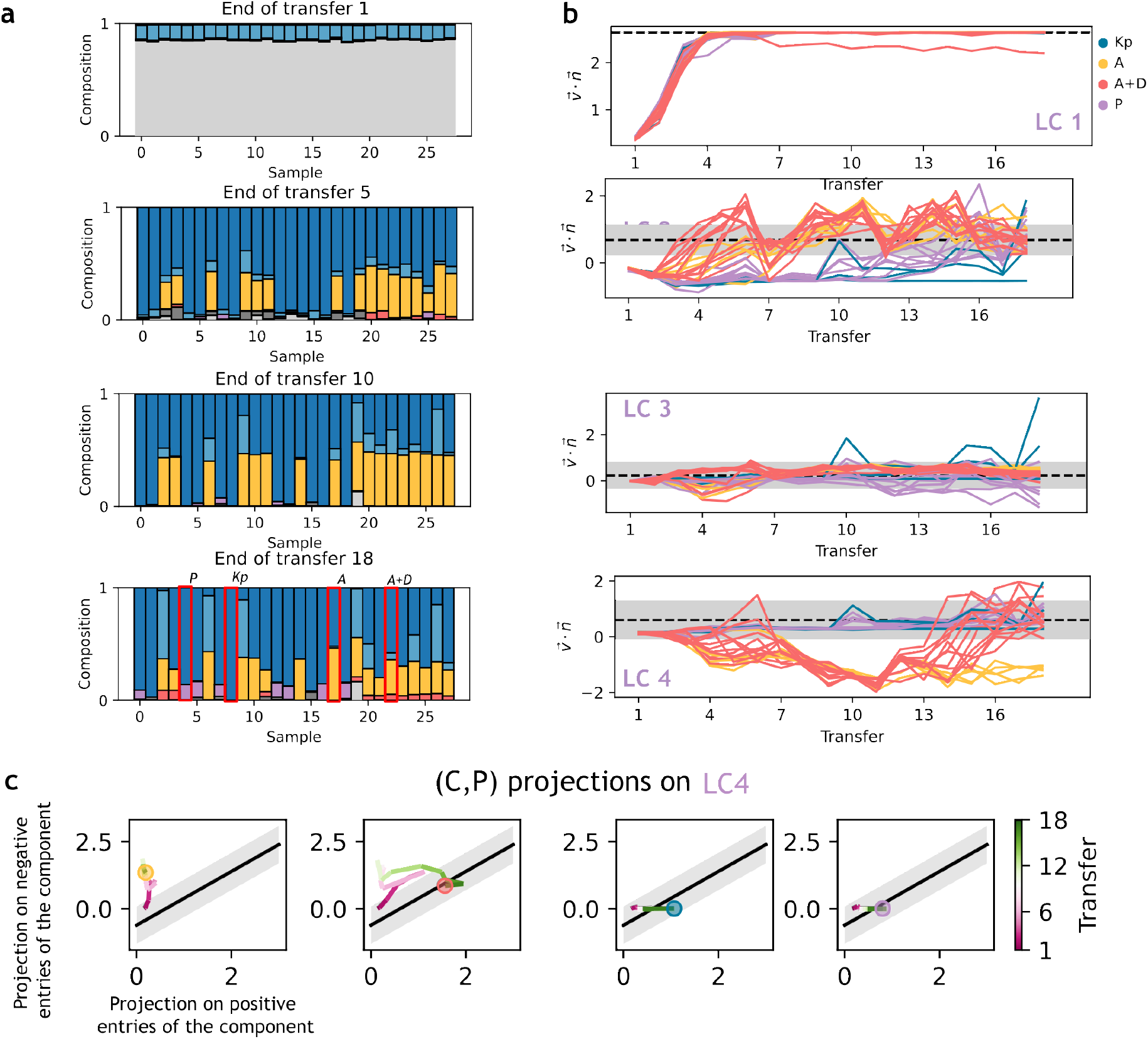
Different low variance modes are fulfilled at different stages of the serial dilution experiment. **(a)** Community compositions at different transfers. Gray indicates taxa not present in the dataset at the final transfer (absent or discarded due to rarity). **(b)** Projection of experimental replicates, at different transfers, on the LCs, showing that different LCs are fulfilled with different timescales. The color of every replicate indicates the macroscopic state of the community after the final transfer, as in Fig. SI 18 (labeled examples in panel A, bottom). The width of the gray areas is 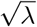. **(c)** Average trajectory, per community state, on the positive and negative projections on LC4 across the 18 transfers of the experiment. Different community types exhibit different trajectories in CP-space, potentially enabling prediction of final community states from states at earlier transfers.

**Figure SI 19.**
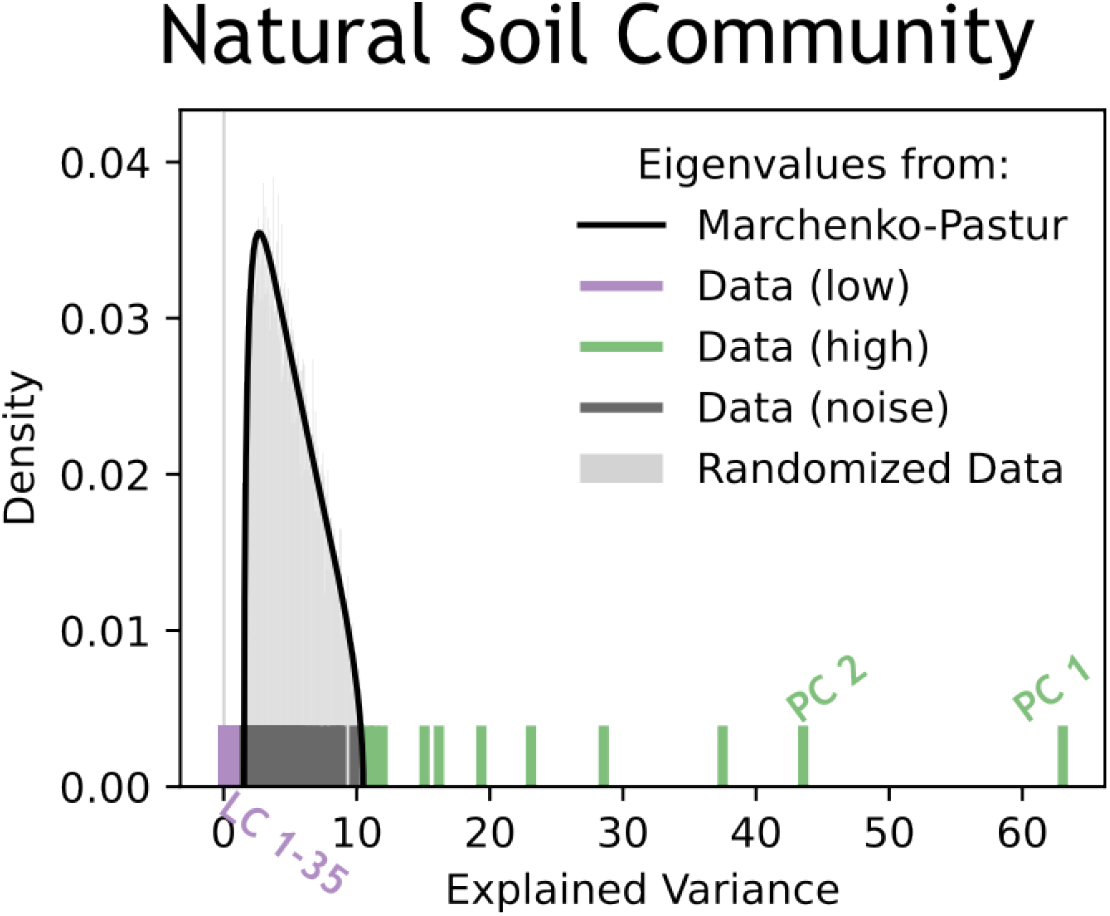
Eigenspectrum of the natural soil community. LCs and PCs are discussed in the main text.

**Figure SI 20.**
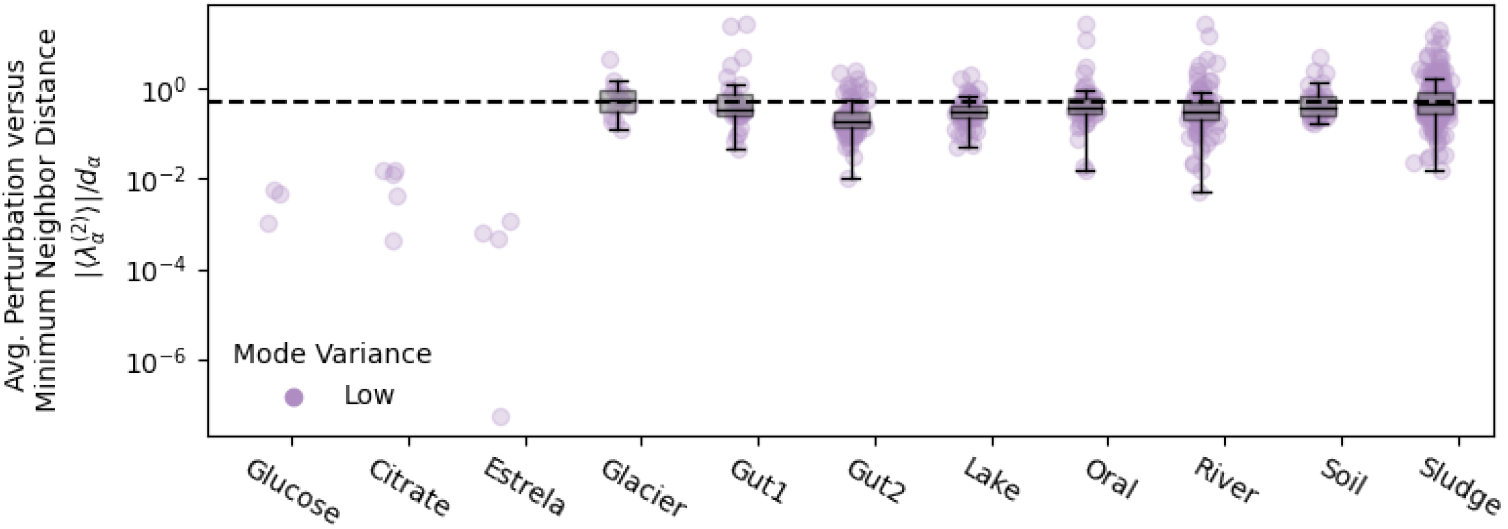
Least components in natural communities are fragile to the addition of new samples. We plot the expected second order correction in the eigenvalue of a least component upon addition of a new sample to the dataset, 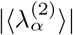 divided by the distance to the nearest neighbor in the eigenvalue spectrum, d_*α*_, for experimental (Glucose and Citrate from [4] and the dataset from Estrela *et al*. [2]) and natural (others) datasets. The dashed line is at 0.5. Across natural datasets, the expected change in eigenvalues is close to the inter-eigenvalue distance, whereas for experimental datasets it is much smaller.

**Figure SI 21.**
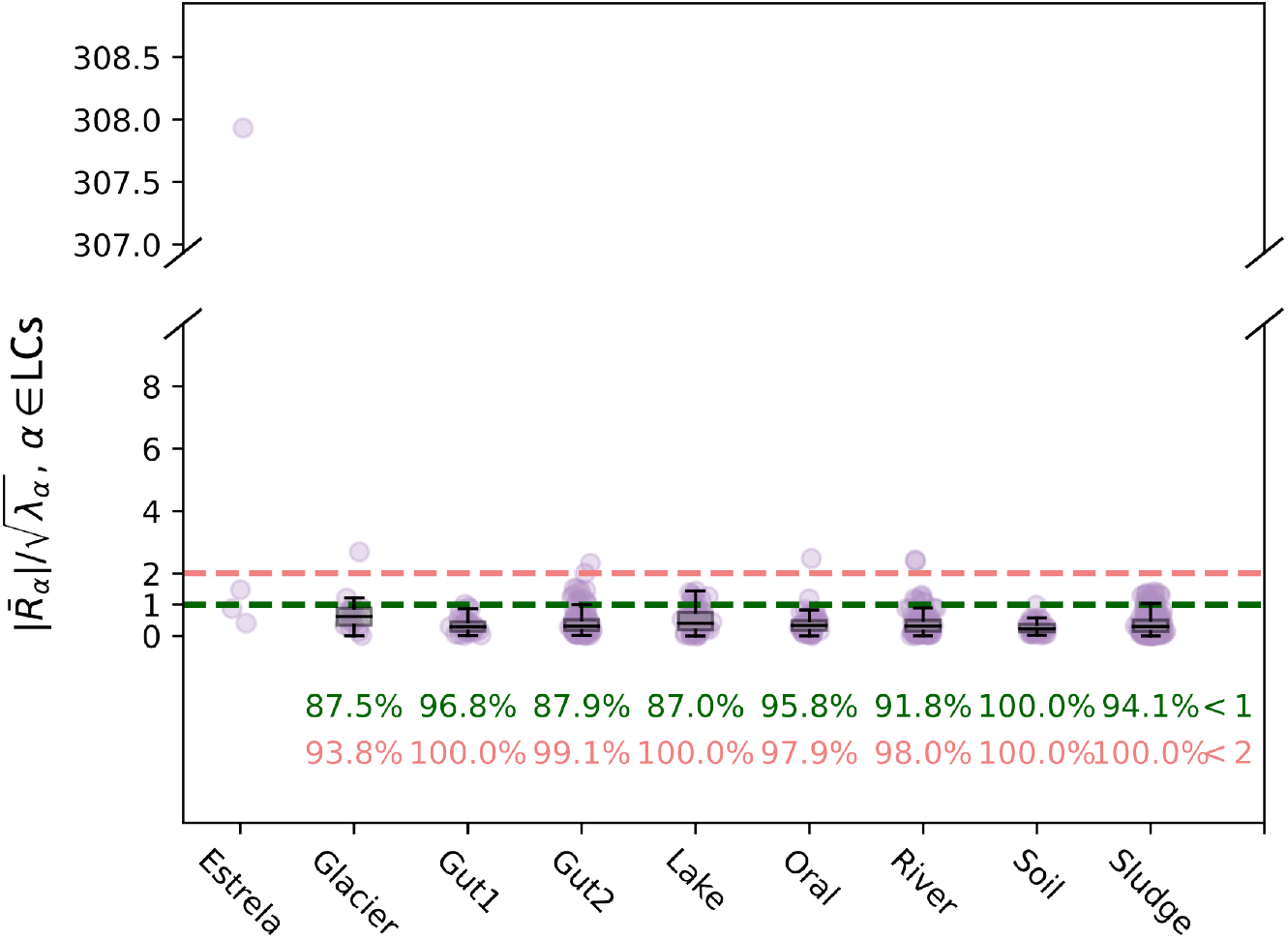
LCs encode for facilitation in natural communities. Across all natural datasets, the average of the projection of the data on the LCs is less than one standard deviation away from 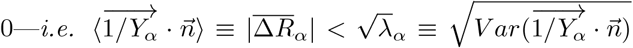 (using the consumer-resource notation of Eq. 1). We also show the experimental dataset of Estrela *et al*. for reference, where all least components infer facilitative interactions, except for the LC of relative abundances (top marker). Numerical values indicate the percentage of LCs with an 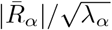 value below 1 (green) and 2 (red).

**Figure SI 22.**
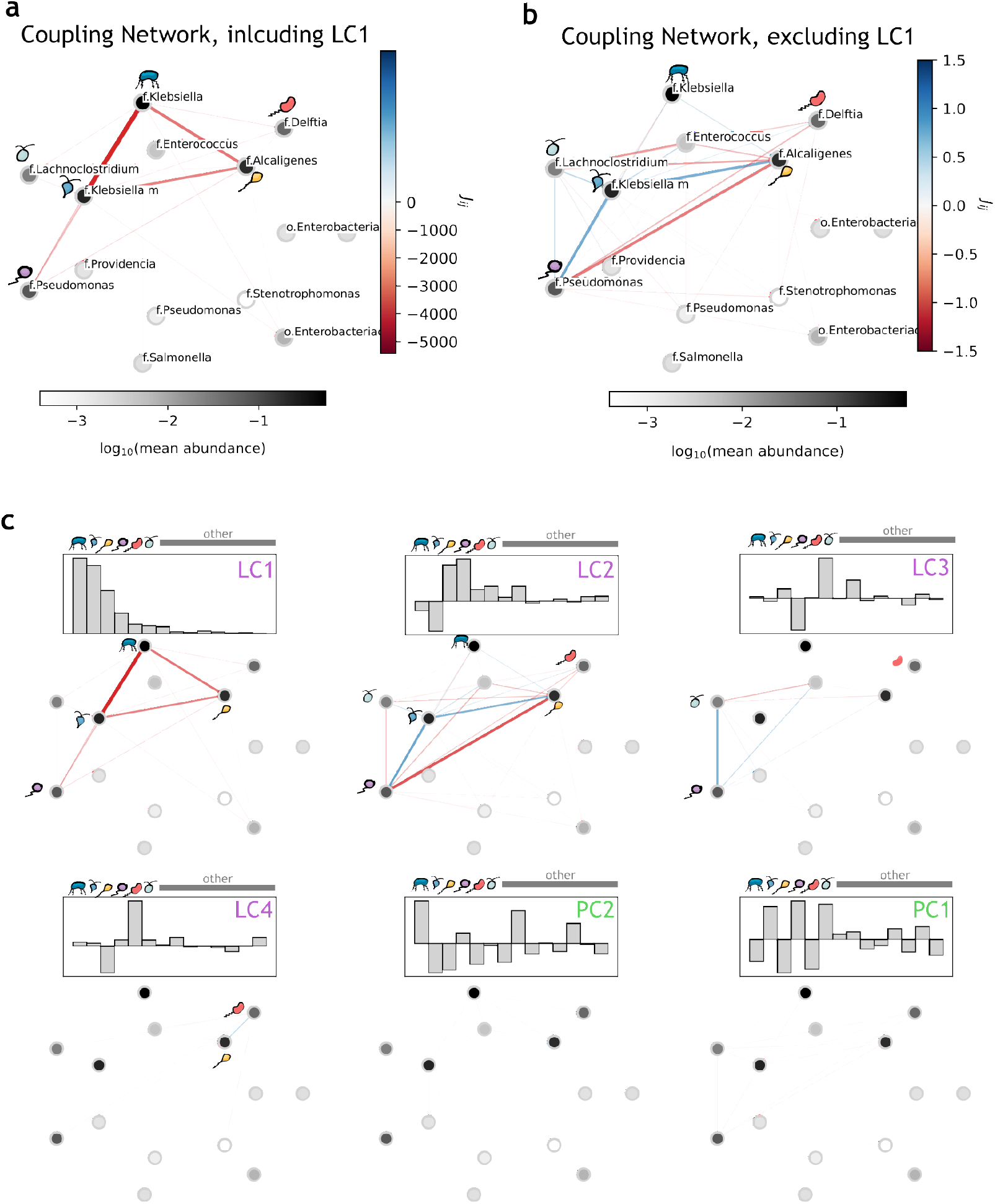
Coupling Network in the Experimental Soil Dataset is dominated by the LC for relative abundances. **(a)** Coupling network (as in Fig. 5) of the taxa in the experimental dataset of Estrela *et al*. [2]. **(b)** Coupling network, without the contribution of LC1 **(c)** Contribution of individual components (entries shown in the gray barplots) to the coupling network. The color code for LC1 is as in panel A. For all other components it’s as in panel B. Bacteria drawings are as in the main text, to ease comparison.

